# Uncovering the link between gut microbiome, highly processed food consumption and diet quality through bioinformatics methods

**DOI:** 10.1101/2023.03.07.530223

**Authors:** Víctor Manuel López Molina, Blanca Lacruz-Pleguezuelos, Enrique Carrillo de Santa Pau, Laura Judith Marcos-Zambrano

## Abstract

The consumption of highly processed foods, along with other dietary and lifestyle poor habits, has an impact on health by increasing the risk of several non-communicable pathologies, such as diabetes. Gut microbiome composition, in specific, can be modulated by nutrients, deriving in different metabolic outcomes that have an influence on this high disease susceptibility and making it a possible therapeutic target for these comorbidities. In this work, gut microbiome of 60 and 46 individuals from 2 different studies focused, among other aspects, on diet-microbiome interactions, was characterised. By means of differential abundance analyses and supervised machine learning techniques based on random forest, gradient boosting and support vector machines, a set of microbial genera that could be potential biomarkers for the differentiation of individuals with poorer dietary patterns was discovered, after comparing coincidences in these taxa among classifiers and testing them for significant differences. Among these, *Dialister*, *Phocea* and *Pseudoflavonifractor* were suggested to have a role in the way highly processed foods affect health negatively, along with *Prevotellaceae NK3B31 group* and an undetermined genus from *Muribaculaceae* in the opposite sense. Furthermore, all the identified genera in this study had already been linked to type 2 diabetes, among which *Bacteroides* and *Pseudoflavonifractor* proved to be differentially abundant in groups of individuals with different levels of biomarkers for this disease. Nevertheless, further research via longitudinal studies and experimental validation of these genera should be carried out to confirm the association of these taxa with diet and diabetes.

## Introduction

Non-communicable diseases (NCDs) are pathologies that are not spread through infection, but are caused by a combination of genetic, physiological, environmental and behavioural factors, such as smoking, unhealthy eating habits, irresponsible alcohol consumption and insufficient physical activity [1]–[3]. The main types of NCDs are cardiovascular diseases (CVDs), cancer, chronic respiratory diseases and diabetes [1]; and they can affect people of all age groups and countries, especially those living in low-and middle-income nations [1]. The relationship between dietary habits and chronic non-communicable diseases, which are responsible for 74% of global deaths yearly [1], has been extensively investigated [4]. A recent systematic analysis of the effects of a suboptimal diet on health estimated that bad dietary habits were responsible for 11 million deaths and 255 million disability adjusted life years (DALYs) in 2017 [4].

Among these poor dietary habits, highly processed food (HPF) consumption is one of the most relevant. The intake of fruits, vegetables and fresh products is quickly decreasing while the consumption of HPFs is increasing worldwide, especially in high-income countries [5]–[8]. In fact, as of 2022, HPFs account for more than 50% of the energy intake in Westernised countries, like the United Kingdom, the United States and Australia [9]–[12]. HPFs are formulations ready for ingestion, made from refined food substances (such as glucose syrup, hydrogenated oils and modified starch) and a combination of simple sugars, salt, fat and multiple additives [5], [9], [10], [13], [14]. As a result, HPFs are rich in refined sugars, fats and salt, while low in proteins, fibres and micronutrients [10], [13].

These foods, which include sugar-sweetened beverages (SSBs), processed meat products, snacks, industrially processed pastries and pre-packaged meals [5], [9], [11], [13], among others, are not recommended for extended consumption, not only for their low nutritional quality, but also their hyper-palatable attributes which promote overconsumption [5], [9], [10], [13], [15]. This induces a lot of economic interest on companies, which make a special effort on the branding and marketing of these products, targeted particularly toward children and young adults [13], contributing on the adoption of unhealthy eating habits by the youngest generations and thereby increasing the risk of developing an NCD. As a matter of fact, several studies have found associations between HPF consumption and NCDs like obesity [16]–[19], cancer [18], [19], CVDs [18]–[23] and metabolic syndrome [18], [19], [24], as well as inflammatory bowel disease (IBD), Crohn’s disease and ulcerative colitis [25].

Moreover, an inverse association between HPF consumption and the Healthy Eating Index (HEI) has been evidenced [26]–[28]. HEI is a measure for the assessment of whether a set of foods aligns with the Dietary Guidelines for Americans (DGA) [29], [30]. It ranges from 0 to 100, with a higher score indicating greater consistency of the diet with the DGA and thus a healthier diet [29]. Therefore, greater highly processed food intake is associated with lower diet quality [28]. On the whole, given the evidence that diet is a risk factor for NCDs, it is imperative to identify healthy dietary patterns to create evidence-based nutritional recommendations and thus prevent the development of these pathologies.

At the same time, there is progressively more evidence about the link between diet and the gut microbiome (GM) [31], [32]. GM comprises the genome of all microorganisms that inhabit the gut and takes part actively in nutritional, metabolic, physiological and immunological processes, as well as in the production of several metabolites [33]. GM has been suggested to be an influential factor in the development of NCDs in interaction with genetic and environmental factors [34], [35], considering that GM participates in processes like energy homeostasis and immunity [34]–[36], apart from evidence of changes in its composition given any of these pathologies or other factors like diet [37].

Taking into account the recent rise in the intake of highly processed foods and a greater incidence of reduced microbial diversity, decreased enzymatic capacity, diminished fermentation and enrichment of microorganisms that degrade gut mucosa [38]–[40], there is a progressively stronger interest in discovering the mechanisms and microbial signatures involved in NCDs [35], [36], [41]–[44], apart from restoring GM by probiotic or prebiotic intake, dietary changes or faecal microbiota transplantation, among other strategies [45]–[47]. Moreover, there is still little consideration of diet-microbiome-host interactions in nutritional recommendations nowadays [48], and there is a widespread lack of awareness in the general public not only about the significant role that GM plays in the progression of NCDs, but also about the remarkable impact that our daily habits have on our health by modulating GM.

Given this situation, the application of computational tools to the analysis of biological data, especially metagenomic data, which are also compositional, heterogeneous and sparse [49], can be highly useful. In fact, these tools offer the possibility of predicting host phenotypes according to taxonomy-based feature selection and disease states, contributing thus to the development of precision medicine. Furthermore, personalised nutrition can be benefitted from the analysis of microbiome data since GM has proven to play a key role in the interindividual variability in nutrition and can be modulated for therapeutic purposes [43], [50]–[57].

Machine Learning (ML), in specific, is already proving to be strongly practical for the analysis of microbiome data, allowing a better understanding of taxonomical and functional diversity within microbial communities. Several models have been developed for the classification of microbial features [58]–[60], the prediction of the development of certain pathologies [43], [61]–[63] and the stratification of patients based on the characterisation of microbial signatures which have previously been linked to certain disease states [64]–[66].

## Motivation and Objectives

Given the evidence regarding the links between GM and diet, and how the latter is a risk factor for NCDs, this work aimed to study whether the relation among HPF consumption, diet quality and NCD risk is mediated by changes in GM composition. On this basis, GM of 60 volunteers, with different HPF intake and HEI, from a 16S ribosomal RNA (rRNA) study was characterised, and then several ML models were built using these data in order to look for potential microbial biomarkers of HPF consumption and unhealthy dietary habits. GM data of 46 volunteers from a whole-genome shotgun sequencing (WGS) study were also used to validate one of these ML models that had previously been trained based on HEI classification of the first cohort.

## Materials and Methods

For this work, two datasets coming from two different projects were taken into consideration. Raw paired-end reads from the variable V3 and V4 regions of the bacterial 16S rRNA gene were collected from a cohort of 70 samples of a 16S rRNA sequencing study developed at the IMDEA Computational Biology group, which will be called “**Study cohort**”, and they were taken into account for the construction of several ML models. Additionally, raw paired-end reads originating from a WGS study carried out in the same group with a cohort of 95 participants, which will be named “**Validation cohort**”, were used for the validation of one of these ML classifiers. Given the differences in the nature of these data, as well as in the way they are analysed, the methods applied to each dataset are exposed in different subsections:

### Data collection and pre-processing

#### Methods for the processing of 16S rRNA sequencing data from the study cohort

Along with the aforementioned raw reads, all necessary metadata for classification were collected. Participants were therefore asked to provide anthropometric and clinical data, information about the consumption frequency of a wide variety of foods, the consumption frequency of HPFs, physical activity frequency and finally fill in a survey regarding the knowledge, related to microbiome and its care, gained after the study.

In order to adapt the metadata for the subsequent metagenomic analyses and ML procedures, pre-processing and homogenisation of these data had to be performed. As for the clinical and anthropometric data questionnaire, in which participants had to indicate, among other information, whether they were taking any medication at the time, how much alcohol they used to drink daily or their family medical history; a standardisation and clustering of these data in simpler groups was carried out.

Regarding the food consumption frequency questionnaire, participants could only reply by using a pre-defined answer, chosen between 9 different options. In order for R to understand these data and simplify them, these answers were renamed as numbers, establishing the following options: 0 (NA), 1 (never or < 1 time per month), 2 (2-4 times per week), 3 (once per day), 4 (1-3 times per month), 5 (5-6 times per week), 6 (2-3 times per day), 7 (once per week), 8 (6 or more times per day) and 9 (4-5 times per day). As for the HPF consumption frequency questionnaire, participants could reply from between 8 different options, which were the same as the previously mentioned ones, except for option 8, which meant “4-6 times per day”.

Finally, as for the physical activity questionnaire, this one was based on the Global Physical Activity Questionnaire (GPAQ) [67], [68], which collects information on physical activity in three settings (activity at work, travel to and from places and recreational activities) as well as sedentary behaviour; and consists in 16 questions (P1-P16). Out of these, 5 questions are replied affirmatively or negatively, while the rest have to be answered using a number of days or hours and minutes. Skips of questions can only be done when any of these 5 questions have been answered negatively. GPAQ relies on metabolic equivalents (METs), which express the intensity of physical activities. One MET is defined as the energy cost of sitting quietly and is equivalent to a caloric consumption of 1 kcal/kg/hour. For GPAQ, 4 METs get assigned to the time spent in moderate activities, and 8 METs to the time spent in vigorous ones. Taking this into account, and based on the answers given by the subjects of this study, three levels of physical activity were established in accordance with GPAQ questionnaire interpretation [67], [68]: low, moderate and high. In order to assign one of these levels to each participant, the total physical activity in MET-minutes/week ([(P2 * P3 * 8) + (P5 * P6 * 4) + (P8 * P9 * 4) + (P11 * P12 * 8) + (P14 * P15* 4)]) was calculated first. Then, the participants were clustered following these criteria:

- High:
  - IF: (P2 + P11) ≥ 3 days AND total physical activity MET minutes per week is ≥ 1500. OR
  - IF: (P2 + P5 + P8 + P11 + P14) ≥ 7 days AND total physical activity MET minutes per week is ≥ 3000.
- Moderate:
  - IF: level of physical activity does not reach criteria for high levels of physical activity AND at least one of the following:
  - IF: (P2 + P11) ≥ 3 days AND ((P2 * P3) + (P11 * P12)) ≥ 3 * 20 minutes. OR
  - IF: (P5 + P8 + P14) ≥ 5 days AND ((P5 * P6) + (P8 * P9) + (P14 * P15)) ≥ 150 minutes. OR
  - IF: (P2 + P5 + P8 + P11 + P14) ≥ 5 days AND total physical activity MET minutes per week ≥ 600.
- Low: IF level of physical activity does not reach the criteria for either high or moderate levels of physical activity.

For their taxonomic assignment, raw paired-end reads were imported into QIIME2 (version 2022.2.0) [69], a platform that integrates multiple tools for the analysis of metagenomic sequences. Data produced or handled by QIIME2, after providing the original data, are stored as artefacts, which present the .qza extension. QIIME2 also generates visualisation data stored as .qzv files. After importing the sequences and considering that demultiplexing was not necessary since a single .fastq file was available for each read, the DADA2 R package (version 1.20) [70], within QIIME2, was used for the subsequent quality control procedure. A filtering process had to be carried out on the artefacts provided by DADA2 to remove microbial taxa associated with mitochondria or chloroplasts. Later on, so as to determine the similarities/dissimilarities between identified microbial features and build a phylogenetic tree, MAFFT [71] and FastTree [72] algorithms were used within QIIME2, respectively. The taxonomic composition of microbial communities was 7 subsequently determined by means of the SILVA database (release 138.1) [73], which assigns taxonomy to features using updated datasets of aligned small (16S/18S, SSU) and large subunit (23S/28S, LSU) ribosomal RNA sequences for all three domains of life.

Once information regarding the identified amplicon sequence variants (ASV) and their absolute abundances, their taxonomy and their associated rooted phylogenetic tree was obtained, all features assigned to the *Eukarya* kingdom and those assigned to none were discarded using the *subset_taxa* function from the phyloseq R package (version 1.38.0) [74]. All abundances were also agglomerated to the genus level using the *tax_glom* function from the same R package, and then transformed into relative abundances by means of the *transform* function from the microbiome R package (version 1.16.0) [75]. Out of 14647 initial taxa, after the aforementioned filtering procedures, 530 features remained.

#### Methods for the processing of WGS data from the validation cohort

Once raw paired-end reads from the 95 samples belonging to the validation cohort were retrieved, clinical, anthropometric, biochemical and metagenomic data were also taken into account and solely from the first visit of the three in which the interventional WGS study to which this cohort belonged consisted in, since only GM data from this visit were available at the time in which this study was carried out. In concrete, among anthropometric data, focus was limited to weight, height and body mass index (BMI) in our work. Values of several dietary components, such as vitamins, minerals and macronutrients were also collected, along with participants’ HEI, even though this information was available only for 46 of the 95 samples of this cohort, as for the first visit of this WGS study. Regarding clinical and biochemical data retrieved in this project, these included all variables typical of a hemogram, along with cholesterol and type 2 diabetes (T2D) biochemical markers’ levels, among others.

In particular, and considering the links among GM, diet and T2D [37], [76], special focus was laid on glycated haemoglobin (HbA1c), a glycaemic trait that, along with fasting glucose (FC), is used to diagnose diabetes [77]. FC, however, was not considered in our work given that the vast majority of participants had normal glucose levels, at least regarding the first visit of this study. Additionally, the homeostatic model 8 assessment method for insulin resistance (HOMA-IR) was taken into account, along with adiponectin (μg/ml). HOMA-IR is a marker of insulin resistance (IR), one of the main pathophysiological mechanisms of diabetes, and is calculated using fasting plasma insulin and fasting plasma glucose [78], [79]. Conversely, adiponectin is a protein involved in fatty acid oxidation and glucose biosynthesis. It is decreased in subjects with insulin resistance and T2D and increased with weight loss and the use of insulin-sensitising drugs [80], [81], exhibiting thus antidiabetic and anti-inflammatory effects, and also acting as an insulin sensitiser.

For the interpretation of these biomarkers’ values, the following cut-offs were established: as for HbA1c (%), normal values are below 5.7, those between 5.7 and 6.4 are associated with prediabetes and those greater than or equal to 6.5 are linked to diabetes, according to [82]. Regarding HOMA-IR, a score below 1.96 means no IR, while a score between 1.96 and 2.99 might indicate IR, which further studies should confirm, and a score greater than or equal to 3 means IR, based on [79]. Finally, as for adiponectin, normal values range from 5 to 30 μg/ml, being those below 5 μg/ml abnormal and possible indicators of IR or other health conditions, in accordance with [83].

First of all, those reads belonging to the same sample were processed together. Read trimming and removal of host reads were then performed using the default configuration of the *read_qc* module from metaWRAP (version 1.3.2) [84]. This pipeline uses Trim Galore to remove Illumina adapters, as well as trim low-quality ends (Phred score < 20) [85]. Subsequently, it removes host contamination by means of BMTagger (version 3.101), for which the hg38 version of the human genome was used [86]. This tool also generates FASTQC quality reports both before and after processing each one of the samples of interest [87].

As for the taxonomic composition of the microbial communities, this was inferred with MetaPhlAn (version 4.0.0) [88], [89], which relies on BowTie2 (version 2.4.5) to map reads against marker genes [90]. All parameters were used with their default values. Given the fact that MetaPhlAn4 relies on a species-level genome bins (SGB) system [89], and this entailed multiple genera not being properly identified, the SGB-based MetaPhlAn4 output was converted into a GTDB-taxonomy-based profile, which rendered more interpretable results. The relative abundances table obtained after this conversion was then agglomerated to the genus level using the *tax_glom* function from the phyloseq R package (version 1.38.0) [74]. Out of 2343 initial taxa, after the aforementioned filtering procedures, 628 features remained.

Once the taxonomic assignment of reads was performed, the Kruskal-Wallis test was additionally used to test differences in the abundance of all identified genera in the validation cohort, grouped in quantiles based on their levels of HbA1c (%), HOMA-IR, adiponectin (μg/ml) and HEI, in order to discover possible associations between GM and these biomarkers related to T2D and IR.

### Classification of participants

For the subsequent diversity analyses, differential abundance analyses and ML classification of volunteers, participants from the study cohort were classified first based on their HPF consumption and then according to their HEI.

Out of the 70 initial volunteers, 10 of them were discarded since there were no data referring neither their HPF consumption frequency nor their HEI. Thereby, as for the first classification, a first group in which all subjects had consumed ≤ 15% of HPFs out of their total food intake (grams/day) and a second group with all participants who had ingested > 15% of HPFs out of their total intake were established, based on the results reported by [91] and also with the objective of getting groups as much balanced in size as possible. These clusters will from this point forward be named “Low HPF consumption” and “High HPF consumption” groups, respectively.

Then, regarding the HEI classification, the following levels were defined: poor (HEI ≤ 50), acceptable (51-60), good (61-70), very good (71-80) and excellent (> 80), based on [92]. However, for the classification of subjects, they were clustered into two groups: a first one with all participants whose HEI was ≥ 61 (good, very good and excellent) and a second one with all volunteers whose HEI was < 61 (acceptable and poor). This classification was performed regardless of HPF consumption frequency or any other criteria, and also in order to balance group sizes as much as possible. These clusters, additionally, will from this point forward be named “Good HEI” and “Poor HEI” groups, respectively.

As for the participants from the validation cohort, they were also classified into two groups, based on their HEI, using the same criteria as those indicated previously and calling them identically. This, however, entailed discarding 47 samples from the initial 95, since their HEI had not been registered, at least regarding the first visit of the study, which was the one considered in our work. Furthermore, 2 more samples had to be dismissed as well, given that they were named identically and there was no way to differentiate them. The final number of samples from this cohort that were classified into two groups was, therefore, 46.

### Microbial diversity and differential abundance analyses

Alpha diversity measurements focus on the differences in the microbial composition within a single sample [93], [94]. For this analysis, functions from the mia R package (version 1.2.7) [95] were used to compute the Chao1 index for richness (function *estimateRichness*) and Shannon and Simpson’s indices for diversity (function *estimateDiversity*) for both cohorts, although the *transform_sample_counts* function from the phyloseq R package (version 1.38.0) [74] was applied first on the validation cohort to transform the relative abundances table to an absolute abundances one. Richness refers to the total number of species in a community, while diversity takes into account species abundances as well [94]. When comparing the Shannon and Simpson’s indices, the first one puts more weight on the richness of a community and less common species, while the latter gives more weight to dominant species [96].

The Wilcoxon rank sum test was used for pairwise tests for both classifications of our cohorts. All *p*-values were also corrected for multiple comparisons using the Benjamini-Hochberg false discovery rate (FDR) correction.

As for beta diversity measures, which were calculated by means of relative abundances of both cohorts, the phyloseq R package (version 1.38.0) [74] was used. These measurements are estimated so as to quantify the differences between communities or samples regarding their composition, and are usually represented as distance matrices [94]. The *distance* function from this package was employed to obtain weighted UniFrac measurements. Additionally, dimensionality reduction of the distance matrix was done by means of principal coordinate analysis (PCoA) via the *ordinate* function. Differences between groups were tested by permutational multivariate analysis of variance (PERMANOVA) with 999 random permutations, using the function *adonis2* from the vegan R package (version 2.6.4) [97].

In addition, statistical differences in microbiome abundances between groups, for both cohorts and their classifications, were tested using the DESeq2 R package (version 1.34.0) [98]. For each cohort, a phyloseq object containing all agglomerated absolute abundances went through an initial filtering process which consisted in removing all features whose total number of counts was less than or equal to 10. A DESeq2 object was created after this for each cohort, and the *DESeq* function was employed to estimate the size factors (using the median of the ratios of observed counts), obtain standard maximum likelihood dispersion estimates and perform differential expression testing by means of a Negative Binomial Wald test, considering the null hypothesis that there is no differential expression across the two groups (log2FC = 0) and assuming a zero-mean normal prior distribution for generalised linear model (GLM) coefficients, in this case, log2 fold changes [99], [100].

Result tables were generated using the *results* function, in which log2 fold changes, *p*-values and adjusted *p*-values were displayed for each genus. This function automatically performs independent filtering based on the mean of normalised counts for each genus, optimising the number of genera which will have an adjusted *p*-value below a given Benjamini-Hochberg FDR cut-off, which was set to 0.05. All log2 fold changes not being between -1 and 1 were discarded in addition to this.

### Machine Learning for classification based on highly processed food consumption and healthy eating index

#### Pre-processing of relative abundances prior to classification

The relative abundances table from the study cohort, containing 530 microbial taxa for both classifications, was subjected to the addition of a pseudo count equal to half the minimal non-zero value in order to remove zero abundance values. A centred log-ratio (CLR) transformation was performed as well using the *transform* function from the microbiome R package (version 1.16.0) [75]. Furthermore, 56 genera had to be discarded since they were all named “uncultured”. Moreover, 274 features with an absolute Pearson correlation coefficient greater than 0.9 were also removed. The number of metagenomic features remaining after these steps was 200, which were the ones used for model construction.

Regarding the validation cohort, the relative abundances table, containing 628 features, was subjected to the same pseudo count and CLR-transformation procedures. However, only those features shared with the study cohort and used for model training were taken into account, considering the validation cohort was indeed used to validate one of the ML models trained with the aforementioned 200 microbial genera.

#### Classifier construction

For the design of ML models, for both HPF and HEI classifications of the study cohort, the caret (version 6.0.93) [101] and mikropml (version 1.4.0) [102] R packages were used. For each one, random forest (RF) [103], gradient boosting (GB) [103] and support vector classifier (SVC) binary models [103] were built, using the randomForest, gbm, xgboost and kernlab implementations available in the caret R package, since mikropml leverages the caret package [102]. Therefore, the total number of built models was 12 and their classification performance was evaluated in each case.

RF and GB are ensemble methods based on decision trees. Ensemble learning methods combine many simple “building block” models with the objective of building a very powerful final model [104]. In the case of RF, trees are grown independently on random samples of the observations and each split on each tree is performed using a random subset of the features [104], [105]. As for GB, given a loss function and a weak learner, this algorithm seeks to find a model that minimises the loss function. The algorithm is initialised with the best guess of the response, the gradient is calculated, and a model is fit to the residuals to minimise the loss function. The current model is added to the previous one and the procedure is repeated a user-specified number of times [106], [107].

As for SVC, it is a hyperplane-based classifier that does not perfectly separate observations in classes, in the interest of greater robustness and better classification of most of the training observations [104].

GM data from the study cohort were randomised into a training set and a test set with an 80/20% split. All models were trained only on microbiome data, and class labels used for prediction were “Low HPF consumption” and “High HPF consumption” for HPF classification and “Poor HEI” and “Good HEI” for HEI classification. All models were trained by leave-one-out-cross-validation (LOOCV) while performing grid search for several hyperparameters depending on the algorithm. Furthermore, no external feature selection procedures were implemented in the design of the models, since the aforementioned algorithms implicitly conduct feature selection and additional procedures are not recommended [101]: in the case of tree-based models, if a predictor is never used in a split, the prediction equation is independent of these data [101], [103], [104], [107]. As for SVCs, they disregard a portion of the training set samples when creating a prediction equation [107]. With relation to tree-based models, the impurity decrease at each split was obtained via the Gini index criterion. The optimal combination of hyperparameters was chosen based on the model’s accuracy.

Finally, labels on the test data were predicted using each one of the aforementioned models, their receiving operating characteristic (ROC) curves were plotted and the area under the curve (AUC) was calculated as a quality measure using the pROC R package (version 1.18.0) [108]. Furthermore, feature importance was estimated for all models employing the *varImp* function from caret, which uses out-of-bag samples for prediction error estimation [101]. As for the models obtained via mikropml, feature importance, which is estimated by permuting each feature 100 times individually and measuring the decrease in the model’s AUC, can also be consulted without using any additional functions [102]. The best model trained on the study cohort’s HEIs was additionally used to predict HEI groups in the validation cohort.

##### Random Forest

Two classifiers, using caret and mikropml, were built for each classification of the study cohort. For the HPF one, grid search was performed for the *mtry* hyperparameter (from 1 to 70) and different values of *ntree* were evaluated (from 1 to 100) for both models. For the HEI classification, values for both hyperparameters were also tested (from 1 to 200 and from 1 to 100, respectively) for the caret-and mikropml-based models.

##### Gradient Boosting

Two classifiers were built for each classification of the study cohort using caret and mikropml. Since the latter only supports the *xgbTree* caret method, the caret model was built using the *gbm* method and the mikropml one was generated using *xgbTree.* For the HPF classification, grid search was performed for the following hyperparameters in the case of the caret-based model: *interaction.depth* (from 1 to 30), *n.trees* (from 1 to 100), *shrinkage* (from 0.1 to 0.9, with intervals of 0.1), *n.minobsinnode* (fixed to 5) and *bag.fraction* (fixed to 0.6). Regarding the mikropml-based classifier, the optimised hyperparameters were these: *nrounds* (from 1 to 50, with intervals of 5), *max_depth* (from 5 to 20), *colsample_bytree* (from 0.1 to 0.9, with intervals of 0.1), *eta* (from 0.1 to 0.5, with intervals of 0.1), *gamma* (fixed to 0), *min_child_weight* (from 1 to 5) and *subsample* (from 0.1 to 0.5, with intervals of 0.1). For the HEI classification, grid search was performed for the following caret-based model’s hyperparameters: *interaction.depth* (from 1 to 49), *n.trees* (from 1 to 30), *shrinkage* (from 0.1 to 0.9, with intervals of 0.1), *n.minobsinnode* (from 2 to 10) and *bag.fraction* (fixed to 0.7). As for the mikropml-based classifier, different values of the following hyperparameters were tested: *nrounds* (from 1 to 25), *max_depth* (from 1 to 25), *colsample_bytree* (from 0.1 to 0.9, with intervals of 0.1), *eta* (fixed to 0.1), *gamma* (fixed to 0), *min_child_weight* (from 1 to 5) and *subsample* (from 0.1 to 0.7, with intervals of 0.1).

##### Support Vector Classifier

Two classifiers were also built for each classification in this case. For the HPF one, and for both caret-and mikropml-based models, grid search was performed for the *C* hyperparameter (from 100 to 2000, with intervals of 5), while the optimal *sigma* hyperparameter value (0.002893997) was calculated using the *ksvm* function from the kernlab R package (version 0.9.31) [109], [110], which estimates this value according to [111]. For the HEI classification, and regarding the caret-based model, the optimal value of *C* was searched from 0.01 to 10, with intervals of 0.1. As for the mikropml-based model, the most proper value of *C* was searched from 0.0001 to 0.1, with intervals of 0.0001. The optimal *sigma* hyperparameter value (0.003362502) was calculated using the *ksvm* function again, and was used for both models.

## Data and code availability

All data and code used to analyse them, as well as to produce our results are available in the following GitHub repository: https://github.com/victor5lm/tfm

## Results

As indicated in the previous section, a study cohort consisting of 60 samples (out of the 70 original ones) and a validation cohort of 46 samples (out of the 95 original ones) were taken into consideration in this study, whose common demographic data can be found in Table 1. Additional information regarding groups from the study cohort made based on their HPF consumption is shown in Table 2, while other data from both cohorts separated into groups according to their HEI can be seen in Table 3.

**Table 1.** Demographic data for both cohorts.

| Variable | Study cohort, N = 60<sup>†</sup> | Validation cohort, N = 46<sup>†</sup> |
| --- | --- | --- |
| **Age** | 35 (30, 48) | 54 (41, 60) |
| **Sex** |  |  |
| Female | 50 (83%) | 31 (67%) |
| Male | 10 (17%) | 15 (33%) |
| **BMI** |  |  |
| Normal | 40 (67%) | 1 (2.2%) |
| Overweight/Obesity | 20 (33%) | 45 (98%) |

<sup>†</sup> Median (IQR); n (%)

**Table 2.** Summary statistics for the study cohort separated into groups based on HPF consumption.

| Variable | Study cohort |  | p-value<sup>2</sup> | q-value<sup>3</sup> |
| --- | --- | --- | --- | --- |
|  | High HPF consumption (> 15 % g/day), N = 33<sup>1</sup> | Low HPF consumption (≤ 15 % g/day), N = 27<sup>1</sup> |  |  |
| **Age** | 33 (26, 46) | 38 (32, 48) | 0.4 | 0.9 |
| **Sex** |  |  | 0.2 | 0.6 |
| Female | 25 (76%) | 25 (93%) |  |  |
| Male | 8 (24%) | 2 (7.4%) |  |  |
| **BMI** |  |  | 0.3 | 0.7 |
| Normal | 20 (61%) | 20 (74%) |  |  |
| Overweight/Obesity | 13 (39%) | 7 (26%) |  |  |
| **Physical activity** |  |  | 0.14 | 0.6 |
| High | 16 (48%) | 8 (30%) |  |  |
| Moderate | 9 (27%) | 14 (52%) |  |  |
| Low | 8 (24%) | 5 (19%) |  |  |
| **Wine consumption<sup>4</sup>** |  |  | >0.9 | >0.9 |
| 1 | 11 (33%) | 10 (37%) |  |  |
| 2 | 4 (12%) | 4 (15%) |  |  |
| 3 | 1 (3.0%) | 0 (0%) |  |  |
| 4 | 11 (33%) | 8 (30%) |  |  |
| 5 | 0 (0%) | 1 (3.7%) |  |  |
| 6 | 1 (3.0%) | 0 (0%) |  |  |
| 7 | 5 (15%) | 4 (15%) |  |  |
| **Beer consumption<sup>4</sup>** |  |  | 0.6 | >0.9 |
| 1 | 8 (24%) | 10 (37%) |  |  |
| 2 | 8 (24%) | 7 (26%) |  |  |
| 3 | 2 (6.1%) | 0 (0%) |  |  |
| 4 | 7 (21%) | 6 (22%) |  |  |
| 5 | 4 (12%) | 2 (7.4%) |  |  |
| 6 | 0 (0%) | 1 (3.7%) |  |  |
| 7 | 4 (12%) | 1 (3.7%) |  |  |
| **Liquor consumption<sup>4</sup>** |  |  | >0.9 | >0.9 |
| 1 | 25 (76%) | 21 (78%) |  |  |
| 2 | 1 (3.0%) | 0 (0%) |  |  |
| 4 | 7 (21%) | 6 (22%) |  |  |
| **Tobacco consumption** |  |  | >0.9 | >0.9 |
| N | 27 (82%) | 22 (81%) |  |  |
| Y | 6 (18%) | 5 (19%) |  |  |

<sup>1</sup> Median (IQR); n (%)<sup>2</sup> Wilcoxon rank sum test; Fisher's exact test; Pearson's Chi-squared test<sup>3</sup> False discovery rate correction for multiple testing<sup>4</sup> Legend:
1: Never or <1 month
2: 2-4 times per week
3: Once per day
4: 1-3 time per month
5: 5-6 times per week
6: 2-3 times per day
7: Once per week

**Table 3.**
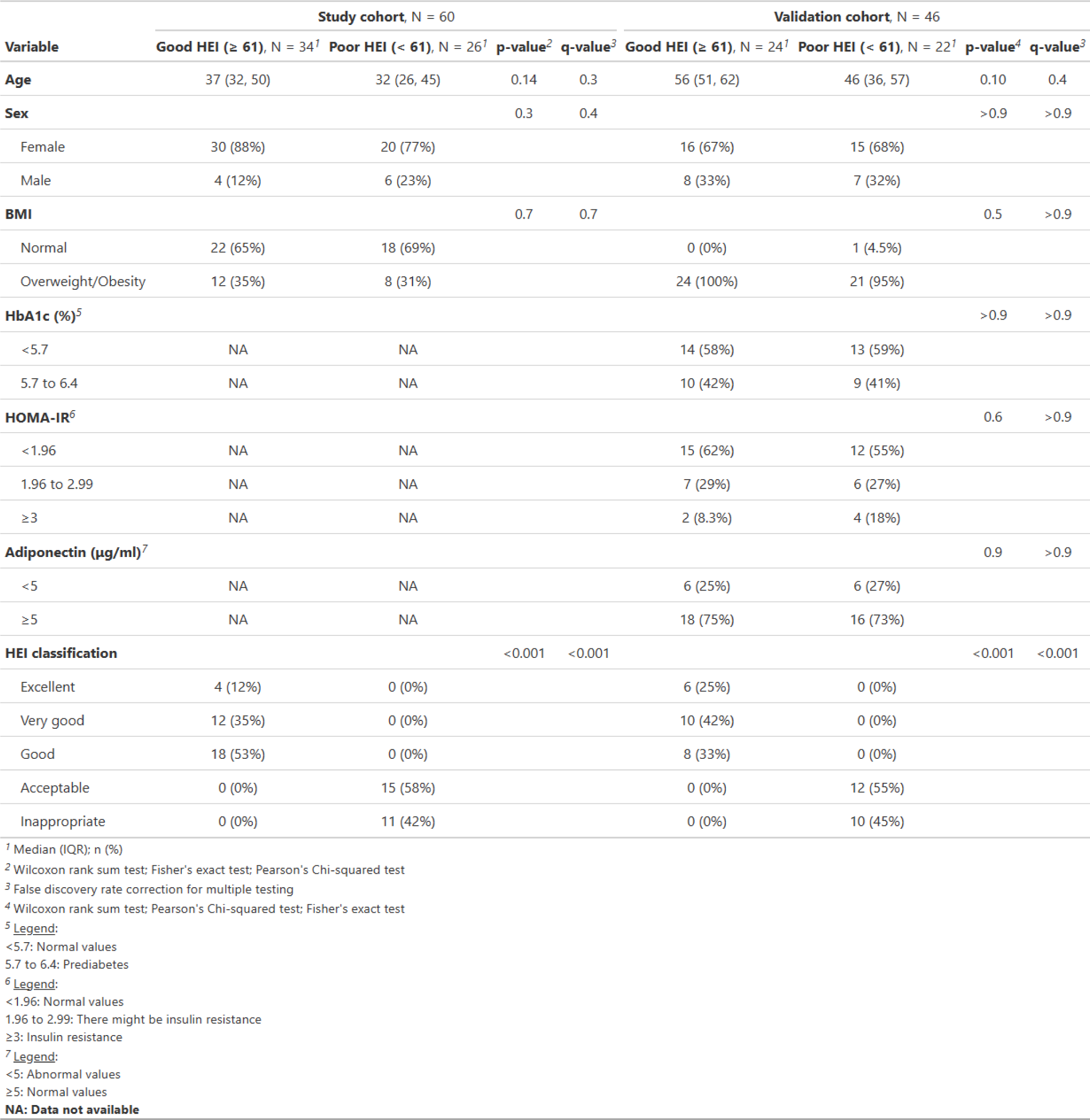
Summary statistics for both cohorts separated into groups based on their HEI.

| Variable | Study cohort, N = 60 |  |  |  | Validation cohort, N = 46 |  |  |  |
| --- | --- | --- | --- | --- | --- | --- | --- | --- |
| | Good HEI ($\geq 61$), N = 34<sup>1</sup> | Poor HEI ($< 61$), N = 26<sup>1</sup> | p-value<sup>2</sup> | q-value<sup>3</sup> | Good HEI ($\geq 61$), N = 24<sup>1</sup> | Poor HEI ($< 61$), N = 22<sup>1</sup> | p-value<sup>4</sup> | q-value<sup>3</sup> |
| **Age** | 37 (32, 50) | 32 (26, 45) | 0.14 | 0.3 | 56 (51, 62) | 46 (36, 57) | 0.10 | 0.4 |
| **Sex** |  |  | 0.3 | 0.4 |  |  | >0.9 | >0.9 |
| Female | 30 (88%) | 20 (77%) |  |  | 16 (67%) | 15 (68%) |  |  |
| Male | 4 (12%) | 6 (23%) |  |  | 8 (33%) | 7 (32%) |  |  |
| **BMI** |  |  | 0.7 | 0.7 |  |  | 0.5 | >0.9 |
| Normal | 22 (65%) | 18 (69%) |  |  | 0 (0%) | 1 (4.5%) |  |  |
| Overweight/Obesity | 12 (35%) | 8 (31%) |  |  | 24 (100%) | 21 (95%) |  |  |
| **HbA1c (%)<sup>5</sup>** |  |  |  |  |  |  | >0.9 | >0.9 |
| <5.7 | NA | NA |  |  | 14 (58%) | 13 (59%) |  |  |
| 5.7 to 6.4 | NA | NA |  |  | 10 (42%) | 9 (41%) |  |  |
| **HOMA-IR<sup>6</sup>** |  |  |  |  |  |  | 0.6 | >0.9 |
| <1.96 | NA | NA |  |  | 15 (62%) | 12 (55%) |  |  |
| 1.96 to 2.99 | NA | NA |  |  | 7 (29%) | 6 (27%) |  |  |
| $\geq 3$ | NA | NA | | | 2 (8.3%) | 4 (18%) | | |
| **Adiponectin ($\mu\text{g/ml}$)<sup>7</sup>** |  |  |  |  |  |  | 0.9 | >0.9 |
| <5 | NA | NA |  |  | 6 (25%) | 6 (27%) |  |  |
| $\geq 5$ | NA | NA | | | 18 (75%) | 16 (73%) | | |
| **HEI classification** |  |  | <0.001 | <0.001 |  |  | <0.001 | <0.001 |
| Excellent | 4 (12%) | 0 (0%) |  |  | 6 (25%) | 0 (0%) |  |  |
| Very good | 12 (35%) | 0 (0%) |  |  | 10 (42%) | 0 (0%) |  |  |
| Good | 18 (53%) | 0 (0%) |  |  | 8 (33%) | 0 (0%) |  |  |
| Acceptable | 0 (0%) | 15 (58%) |  |  | 0 (0%) | 12 (55%) |  |  |
| Inappropriate | 0 (0%) | 11 (42%) |  |  | 0 (0%) | 10 (45%) |  |  |

<sup>1</sup> Median (IQR); n (%)<sup>2</sup> Wilcoxon rank sum test; Fisher's exact test; Pearson's Chi-squared test<sup>3</sup> False discovery rate correction for multiple testing<sup>4</sup> Wilcoxon rank sum test; Pearson's Chi-squared test; Fisher's exact test<sup>5</sup> Legend:
<5.7: Normal values
5.7 to 6.4: Prediabetes
<sup>6</sup> Legend:
<1.96: Normal values
1.96 to 2.99: There might be insulin resistance
$\geq 3$ : Insulin resistance<sup>7</sup> Legend:
<5: Abnormal values
$\geq 5$ : Normal values**NA: Data not available**

Samples from the study cohort had been first divided into two groups, based on the results reported by [91], according to their HPF consumption, depending on whether this was low (≤ 15%) or high (> 15%). Furthermore, both cohorts had been clustered into two groups according to their HEI, depending on whether this was good (≥ 61) or poor (< 61) and considering the HEI levels detailed in [92].

When testing for significant differences between groups in the variables shown in Tables 2 and 3, including alcohol consumption, smoking, physical activity and some glycaemic traits, these were not found neither for any cohort nor any classification, as assessed by Wilcoxon rank sum test for continuous variables, and Pearson’s Chi-squared and Fisher’s exact tests for categorical ones, and assuming the data do not fit a normal distribution, which is characteristic of microbiome data [112]. This lack of differences justifies why our ML models were trained, for both classifications of our study cohort, only on microbiome data.

### Classification based on highly processed food consumption

When classifying our study cohort according to their HPF consumption, the following results were obtained regarding diversity measures, differential abundance analysis between groups and performance of the most precise ML classifier.

#### Diversity measurements

As for this cohort, alpha diversity was first evaluated in order to look for possible significant differences between groups in GM composition, given previous evidence of alterations in GM diversity being linked to metabolic imbalance and metabolic diseases [113]. However, the analysis of GM richness and diversity between subjects who had a lower HPF consumption out of their total food intake (grams/day) and a higher consumption showed no significant differences in the whole population for any of the tested indices (Figure 1).

**Figure 1.**
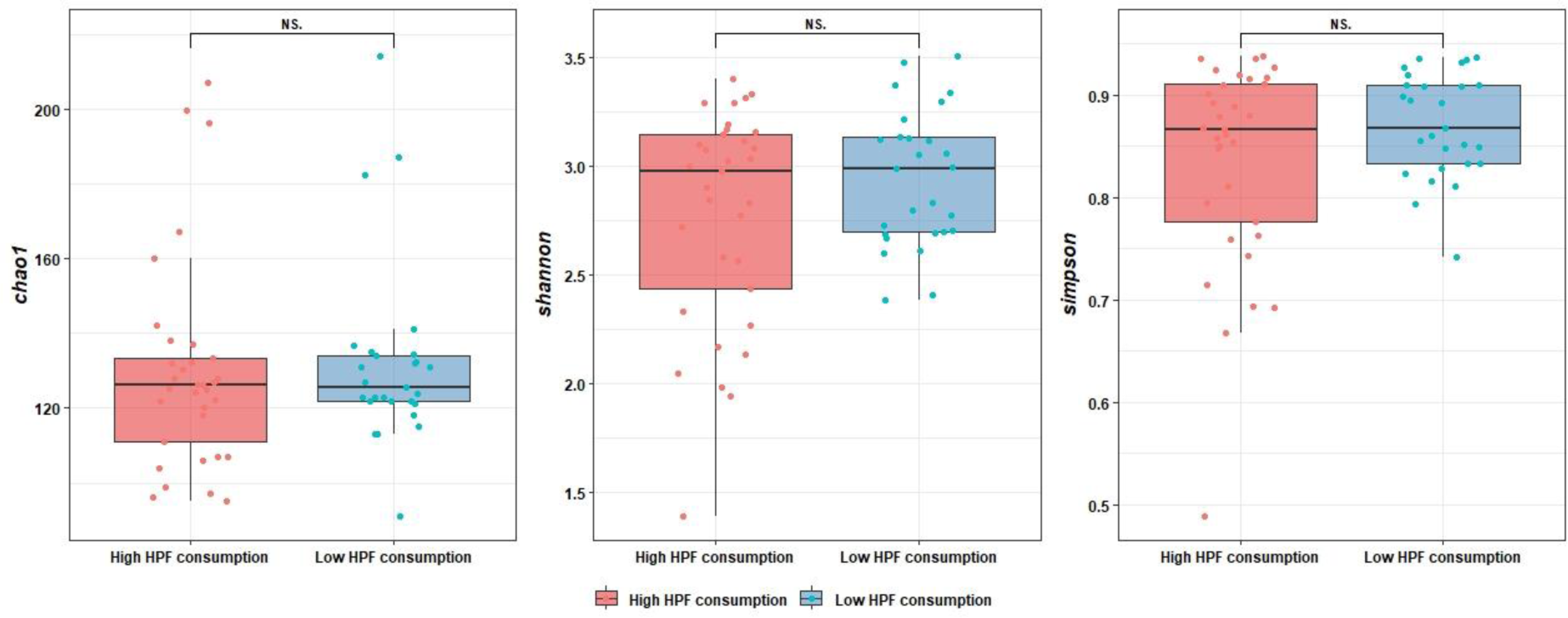
Alpha diversity differences between groups. No significant differences were found in the Wilcoxon rank sum test for the Shannon and Simpson diversity indices (*p* = 0.626 and *p* = 0.626 respectively), as well as the Chao1 richness index (*p* = 0.742). NS.: non-significant.

Additionally, with the objective of evaluating differences among samples as for their community composition, beta diversity was also calculated using the weighted UniFrac index. For our study cohort, no significant differences were found between groups, as assessed by PERMANOVA, and the explained variance was very low (1.14%). As a result, groups could not be separated properly by PCoA, as can be seen in Figure 2.

**Figure 2.**
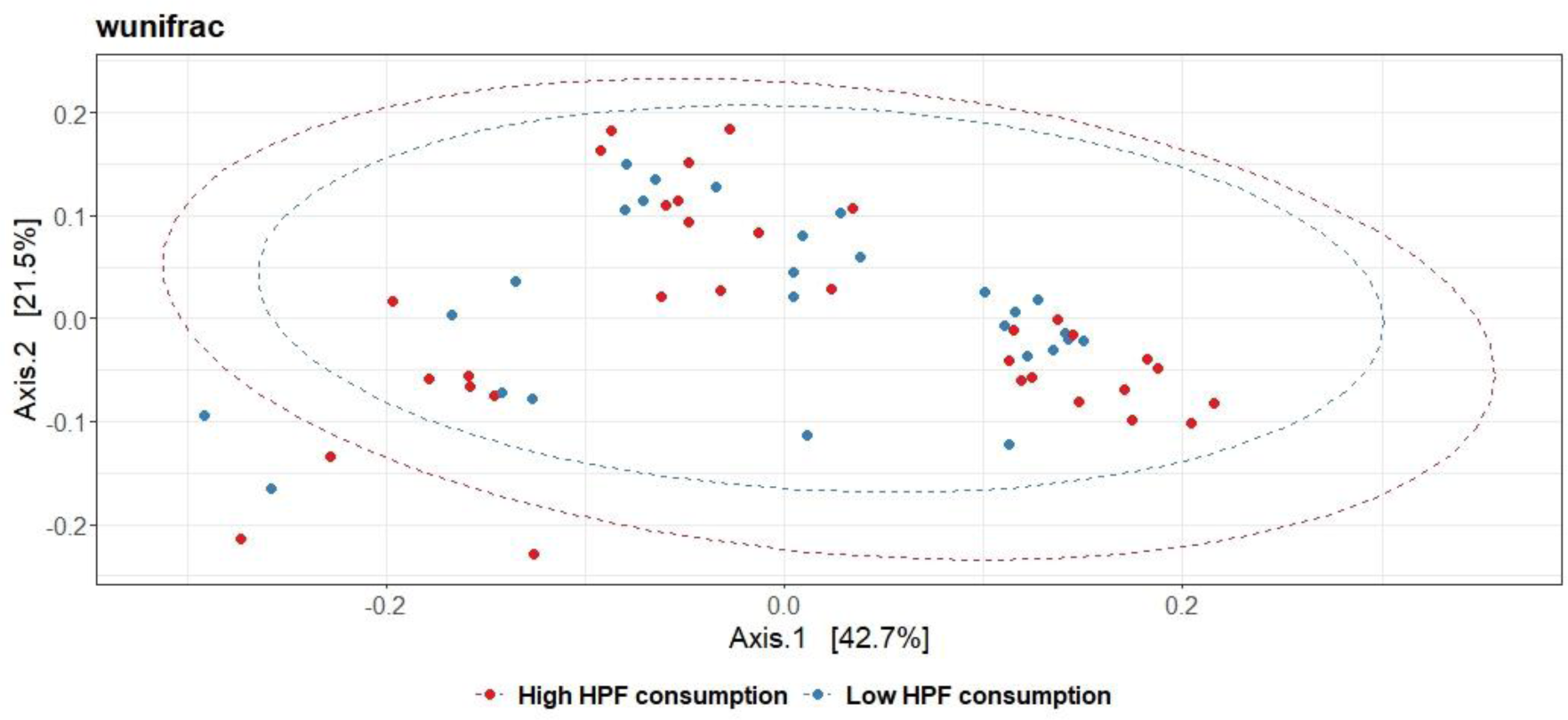
Beta diversity analysis for the study cohort classified based on their HPF consumption: PCoA plot. Weighted UniFrac measurement (*wunifrac*) was calculated with phyloseq and represented via PCoA. Differences between groups were tested with PERMANOVA with 999 permutations. Ellipses represent 95% confidence intervals.

#### Differential abundance analysis

GM profile between the population whose HPF intake was low (≤ 15%) and high (> 15%) was compared by using DESeq2. Without considering adjusted-FDR *p*-values, the participants with low HPF consumption presented significantly higher abundances of the following genera: *Prevotellaceae NK3B31 group*, *Rikenellaceae RC9 gut group*, *Muribaculum*, an undetermined genus from the *Muribaculaceae* family and *Anaeroplasma*. Out of these, the first four belong to the *Bacteroidetes* phylum, while *Anaeroplasma* pertains to *Firmicutes*. On the other hand, the *Dialister* (*Firmicutes* phylum), *Rothia* (*Actinobacteria* phylum) and *Methanobrevibacter* (*Euryarchaeota* phylum) genera were underrepresented in the subjects whose HPF consumption was low. However, when adjusting *p*-values, significant differences were found only for *Prevotellaceae NK3B31 group*, as shown in Table 4.

**Table 4.** Bacterial genera (p < 0.05) analysed by DESeq2 between subjects from the study cohort who had a low (≤ 15%) and high (> 15%) consumption of HPFs.

| Bacterial name | log2FC | *p*-value | FDR-adj. *p*-value |
| --- | --- | --- | --- |
| *Prevotellaceae*<br>*NK3B31* group | 2.685 | $2.766 \times 10^{-5}$ | **$9.1 \times 10^{-3}$** |
| *Rikenellaceae* RC9<br>gut group | 2.012 | $3.667 \times 10^{-4}$ | 0.060 |
| *Muribaculaceae*<br>(undefined genus) | 1.408 | 0.003 | 0.294 |
| *Dialister* | -1.224 | 0.009 | 0.613 |
| *Methanobrevibacter* | -4.568 | 0.016 | 0.762 |
| *Muribaculum* | 1.939 | 0.044 | 0.916 |
| *Anaeroplasma* | 1.776 | 0.043 | 0.916 |
| *Rothia* | -2.540 | 0.046 | 0.916 |

Significant values in bold type.

#### Machine Learning for classification of the study cohort

For this study cohort, 3 binary ML models were built using the caret R package and other 3 by means of the mikropml R package for this HPF-based classification. As for the caret models, RF, stochastic GB and Radial Basis Function (RBF) Kernel SVC algorithms were employed, which were trained exclusively on the CLR-transformed relative abundances table and tested against the true classification to the two categories of the HPF classification using LOOCV. The same cross-validation (CV) procedure and algorithms were used for the mikropml-based ML models, except for the stochastic GB one, which was substituted for an extreme GB algorithm, which is a more regularised version of the first one [103].

All models were trained splitting the data and the accuracy reached during CV, as well as the AUC over the test set, were used as model quality measures. Different hyperparameter values, depending on the algorithm, were tested and the combination reaching the highest accuracy during CV for each model was chosen. The resulting classifiers were used to predict class labels on the test set. The most precise model for this HPF-based classification will be shown next, while the performance metrics of the rest of the models are shown in Supplementary Table 1.

Out of the 6 models built for the HPF classification of our study cohort, which yielded in general adequate results (AUCs between 0.63 and 0.9), the one that achieved the highest AUC (0.9) over the test set was obtained using the mikropml R package with the RF algorithm, where *ntree* = 17 and *mtry* = 15. Thus, this model satisfactorily classified participants in their corresponding group using only CLR-transformed GM data. Then, the ROC curve for this binary model and for our other models were plotted (Fig. 3 and Supplementary Figure 1), as well as variable importance plots (Supp. Fig. 2) for both this model and the other 5 ones. In conclusion, our model allowed us to classify volunteers according to their HPF consumption.

**Figure 3.**
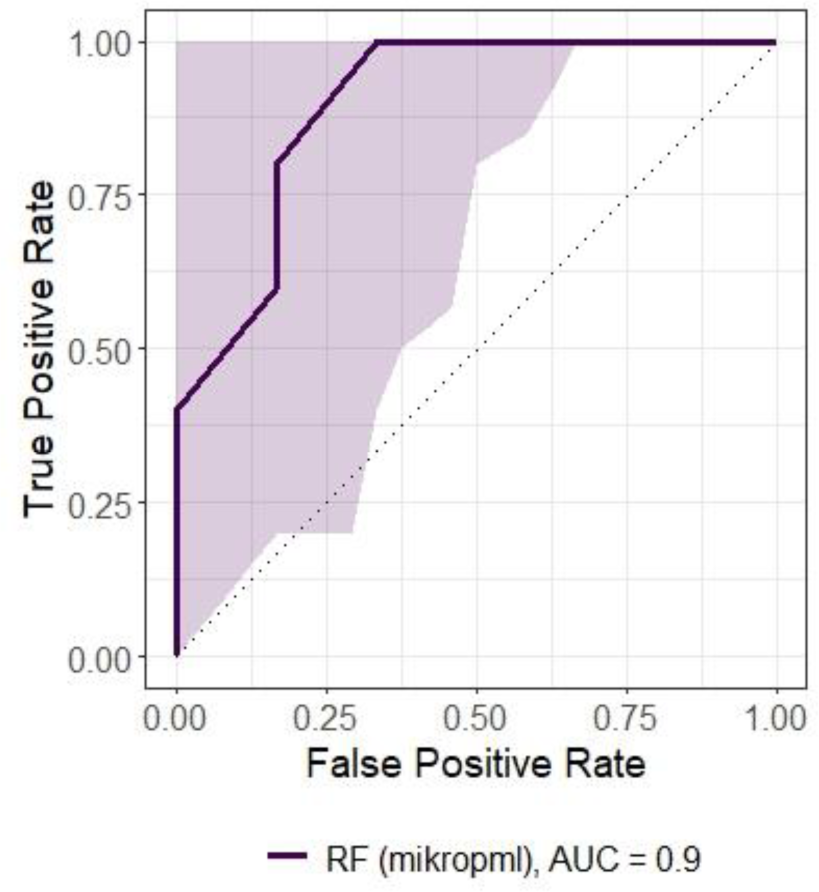
ROC curve for the RF model trained on the CLR-transformed relative abundances table according to the HPF-based classification. Shadowed areas represent 95% confidence intervals for sensitivity.

As for biomarker search, this was performed by comparing the 20 most important features in this model with the ones from the rest of the ML models built for this classification (Supp. Table 2), and testing them for differences in CLR-transformed abundances between both HPF groups (Wilcoxon rank sum test). The following genera were detected as potential biomarkers in the low HPF consumption group: *Prevotellaceae NK3B31 group* (*p =* 0.009, 3 classifiers agreed on its importance), an undefined genus from *Muribaculaceae* (*p =* 0.015, 2 classifiers agreed on its importance) and *Moryella* (*p* = 0.015, 2 classifiers agreed on its importance). As for those volunteers with a higher HPF consumption, *Phocea* (*p* = 0.04, indicated by our caret-based SVC model to be important), was also found to be a possible biomarker, as shown in Supp. Fig. 3. Some of these genera were selected to be important according to our best classifier (Figure 4), while others were not chosen by this one in particular but did be chosen by others, as stated before (Supp. Fig. 2).

**Figure 4.**
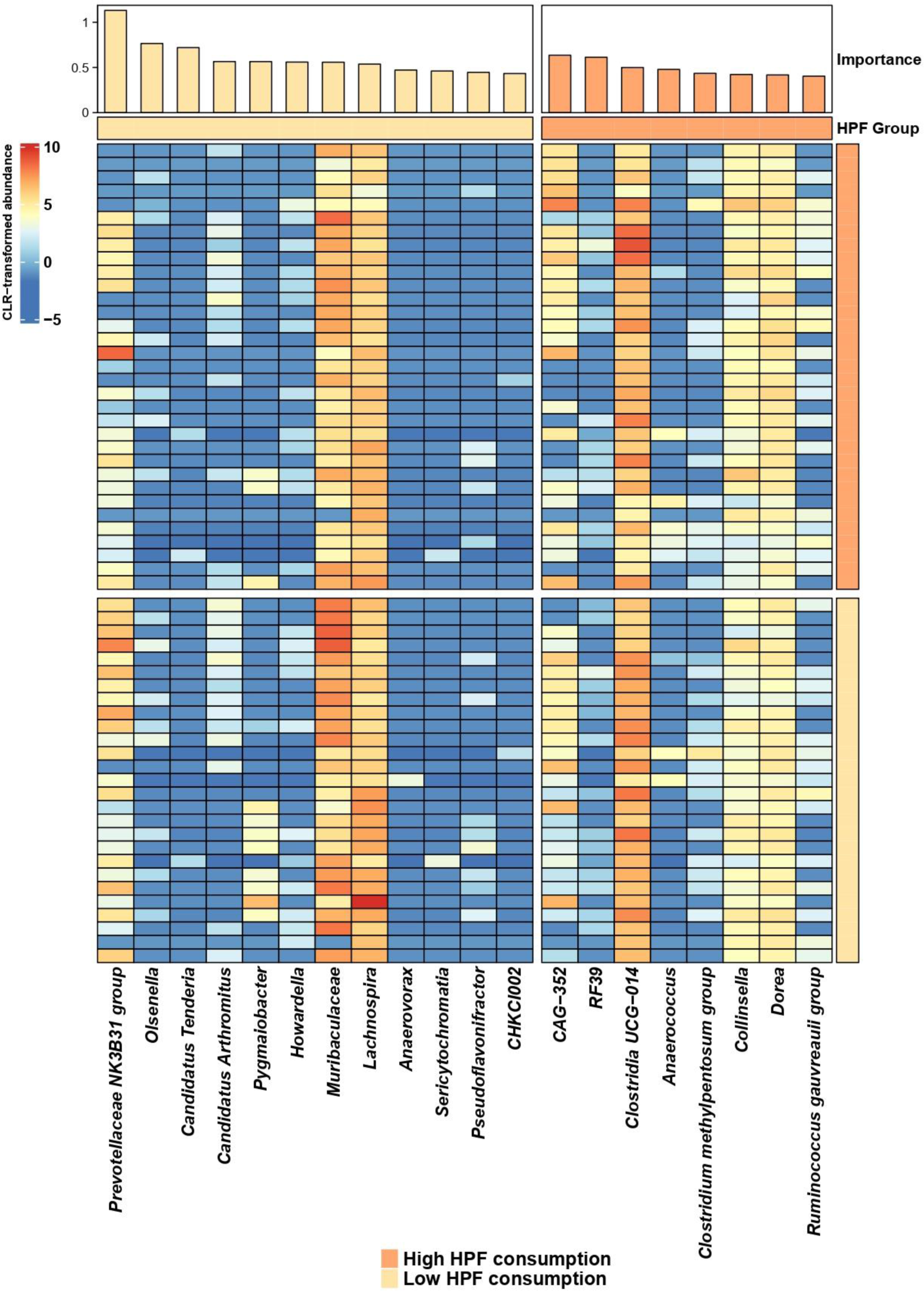
20 most important features according to the RF model based on the HPF classification. Feature importances are plotted at the top of the figure. Bar height indicates feature importance and bar colour indicates whether the genus is enriched in the low consumption group (light orange) or the high consumption one (dark orange). Taxa are sorted according to the group where they are enriched and in descendent order. A heatmap of CLR-transformed relative abundances is plotted below the bar plot. The bar that separates both plots and the one on the right indicate the group in which each genus is more abundant and the one the participants belong to, respectively.

### Classification based on the Healthy Eating Index

When classifying both cohorts according to their HEI, the following results were obtained regarding diversity measures and differential abundance analysis between groups. The best ML classifier’s performance, trained on GM data from our study cohort, will also be shown in this subsection, along with its validation using our validation cohort.

#### Diversity measurements

For both cohorts, alpha diversity was first evaluated for the same purposes as those detailed in section 4.1.1. However, the analysis of GM richness and diversity between subjects who had a good HEI (≥ 61) and a poor HEI (< 61) showed no significant differences neither for any of the tested indices nor for any of our study (Supp. Fig. 4) and validation (Supp. Fig. 5) cohorts. Similarly, when evaluating beta diversity using the weighted UniFrac index, no significant differences were found between groups, as assessed by PERMANOVA, for neither our study (Supp. Fig. 6) nor validation (Supp. Fig. 7) cohorts.

#### Differential abundance analysis

##### Study cohort

GM profile between the population whose HEI was good and poor was compared by using DESeq2. Without considering adjusted-FDR *p*-values, the participants whose HEI was good presented significantly higher abundances of: *Gastranaerophilales* (*Cyanobacteria* phylum), *Prevotella* (*Bacteroidetes* phylum) and *Intestinimonas* (*Firmicutes* phylum). On the other hand, the *Erysipelotrichaceae UCG-003* and *Anaeroplasma* genera, which belong to the *Firmicutes* phylum, were underrepresented in the subjects with good HEI. However, when adjusting *p*-values, significant differences were found only for *Gastranaerophilales*, as shown in Table 5.

**Table 5.** Bacterial genera (*p* < 0.05) analysed by DESeq2 between subjects from the study cohort who had a good (≥ 61) and poor (< 61) HEI.

| Bacterial name | log2FC | *p*-value | FDR-adj. *p*-value |
| --- | --- | --- | --- |
| *Gastranaerophilales* | 3.098 | $1.722 \times 10^{-5}$ | **0.008** |
| *Erysipelotrichaceae UCG-003* | -1.339 | $2.346 \times 10^{-4}$ | 0.054 |
| *Prevotella* | 1.407 | $1.091 \times 10^{-2}$ | 0.997 |
| *Intestinimonas* | 1.064 | $1.637 \times 10^{-2}$ | 0.997 |
| *Anaeroplasma* | -2.149 | $1.800 \times 10^{-2}$ | 0.997 |

##### Validation cohort

GM profile between the population whose HEI was good and poor was compared by using DESeq2. Without considering adjusted-FDR *p*-values, the participants whose HEI was good presented significantly higher abundances of the following genera: *Duodenibacillus* (*Proteobacteria* phylum), *Akkermansia* (*Verrucomicrobiota* phylum), *CAG-353, CAG-115, Lachnospira* and *Acetatifactor* (all of these belonging to the *Firmicutes* phylum). No genera were, nonetheless, underrepresented in the subjects with good HEI. Furthermore, when adjusting *p*-values, no significant differences were found for any genera, as shown in Table 6.

**Table 6.**
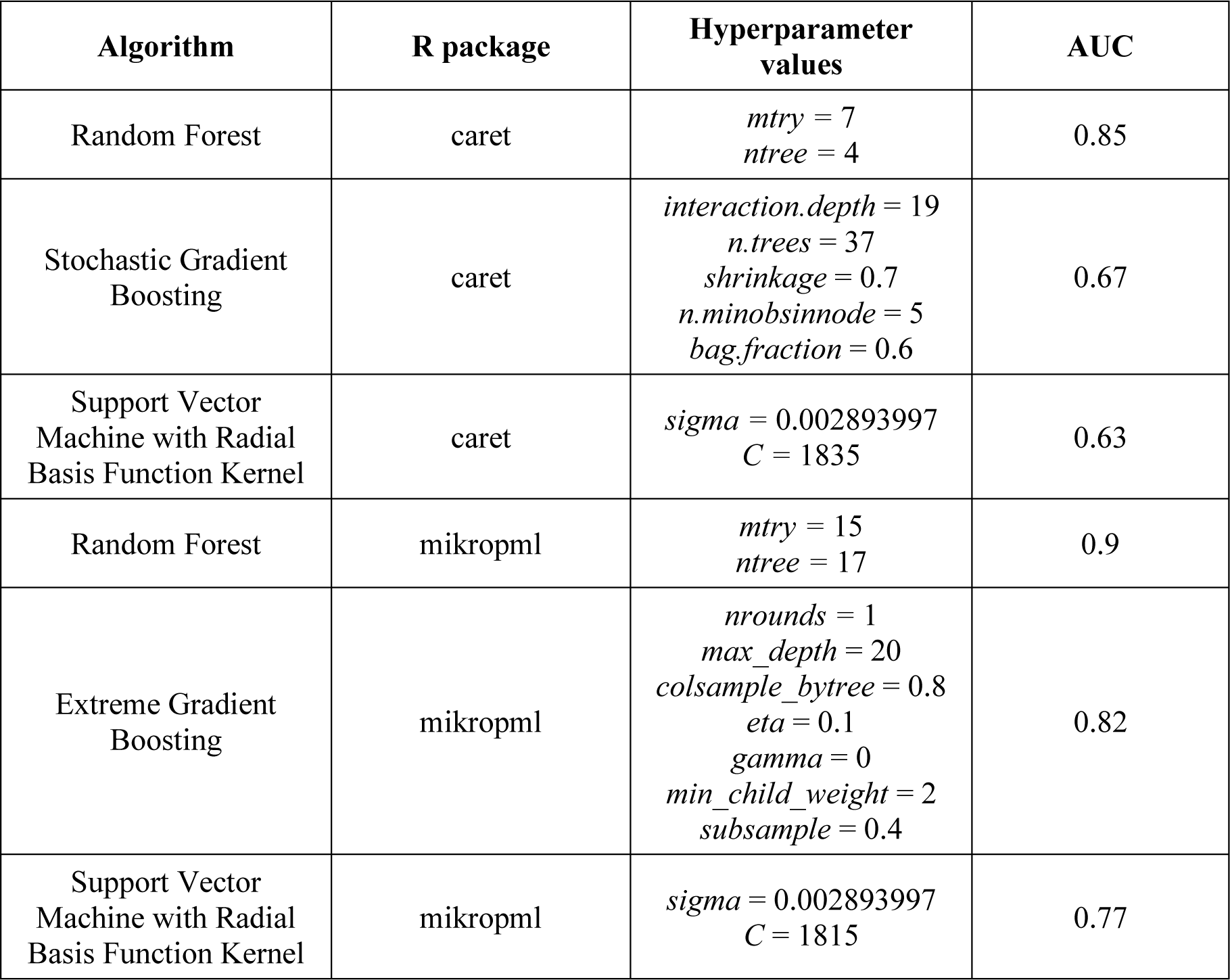
Bacterial genera (p < 0.05) analysed by DESeq2 between subjects from the validation cohort who had a good (≥ 61) and poor (< 61) HEI.

#### Machine Learning for classification of the study cohort

For our study cohort, 6 additional binary ML models were built using the caret and mikropml R packages for the HEI-based classification. As for the caret models, RF, stochastic GB and RBF-Kernel SVC algorithms were employed, which were trained exclusively on the CLR-transformed relative abundances table and tested against the true classification to the two categories of the HEI classifications using LOOCV. The same CV procedure and algorithms were used for the mikropml-based ML models, except for the stochastic GB one, which was substituted for an extreme GB algorithm.

All models were trained splitting the data and the accuracy reached during CV, as well as the AUC over the test set, were used as model quality measures. Different hyperparameter values, depending on the algorithm, were tested and the combination reaching the highest accuracy for each model was chosen. The resulting models were used to predict class labels on the test set. The most precise model for the HEI-based classification will be shown next, while the performance metrics of the rest of the built classifiers are shown in Supplementary Table 3.

Out of the 6 models built for the HEI classification of our study cohort, which yielded in general fair results (AUCs between 0.57 and 0.85), the model that achieved the highest AUC (0.85) over the test set was obtained with mikropml and the RF algorithm, where *ntree* = 88 and *mtry* = 64. Thus, this model satisfactorily classified participants in their corresponding group using only CLR-transformed GM data. Then, the ROC curve for this binary model and for our other models (Fig. 5 and Supp. Fig. 8) were plotted, as well as variable importance plots (Supp. Fig. 9) for all classifiers. In conclusion, our model allowed us to classify volunteers according to their HEI. Additionally, this classifier was validated employing the CLR-transformed relative abundances table from our validation cohort. Prediction on this dataset yielded an AUC of 0.7 (Figure 6).

**Figure 5.**
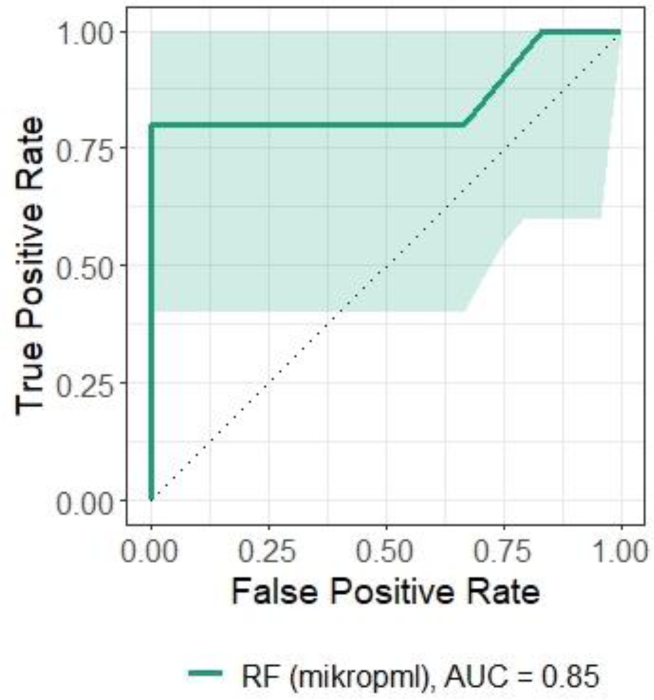
ROC curve for the RF model trained on the CLR-transformed relative abundances table according to the HEI classification. Shadowed areas represent 95% confidence intervals for sensitivity.

**Figure 6.**
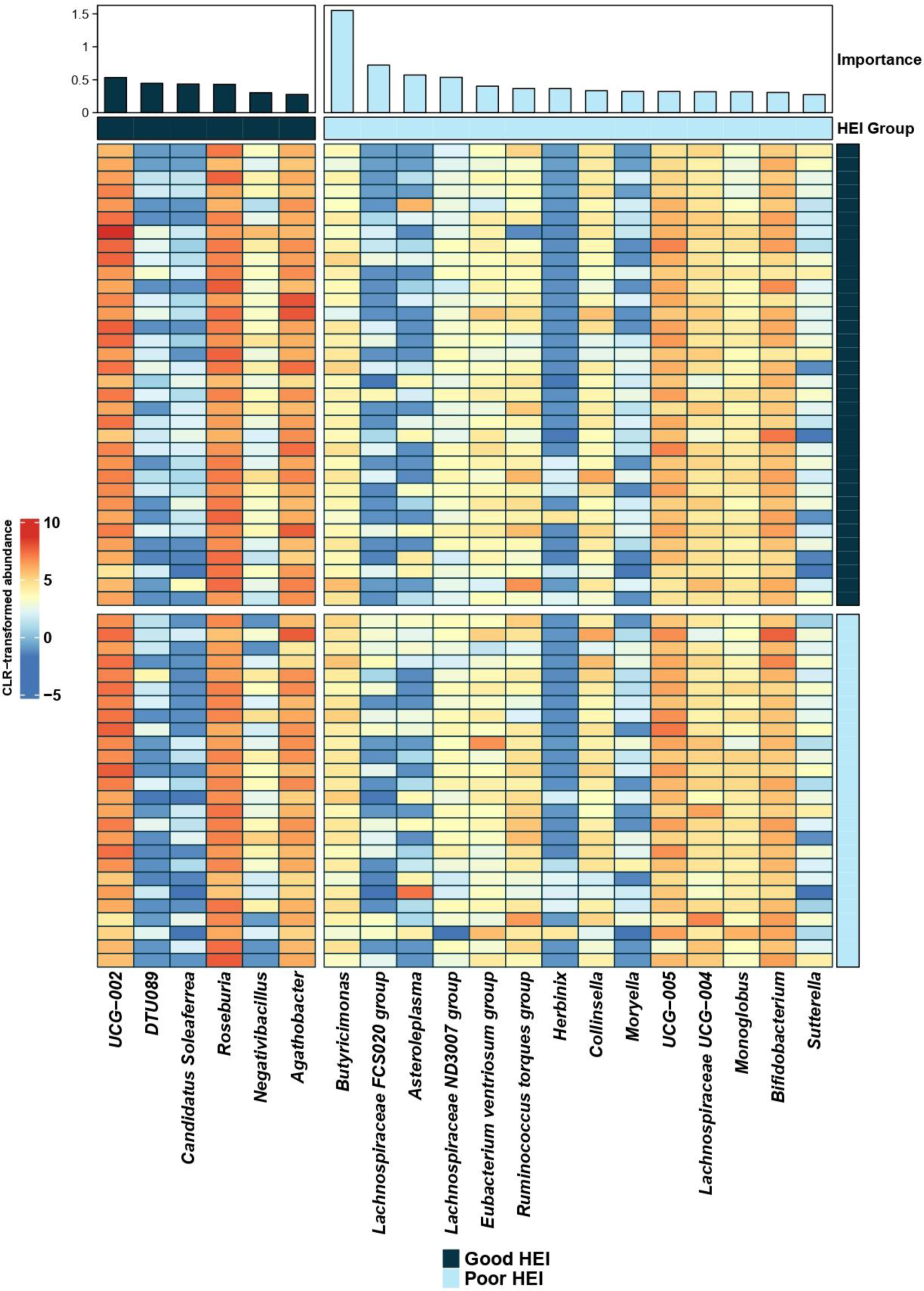
ROC curve for the validation cohort using the RF model trained on the CLR-transformed relative abundances table (HEI classification). Shadowed areas represent 95% confidence intervals for sensitivity.

Biomarker search was performed comparing the 20 most important features in this RF model with the ones from the rest of our classifiers (Supp. Table 4), and testing them for differences in CLR-transformed abundances between both HEI groups (Wilcoxon rank sum test). The following genera were detected as potential biomarkers in participants from our study cohort whose HEI was good: *Candidatus Soleaferrea* (*p =* 0.023, 2 classifiers agreed on its importance)*, Intestinimonas* (*p =* 0.033, highlighted by DESeq2) and *CAG-352* (*p* = 0.048, important according to our caret-based GB model). As for volunteers with poor HEI, *Butyricimonas* (*p* = 0.033, 4 classifiers agreed on its importance) was found to be a possible biomarker as well (Supp. Fig. 10). *Lachnospira* and *CAG-115*, highlighted by DESeq2, were also detected as marker candidates in the validation cohort (Supp. Fig. 11). Some of these genera were selected to be important by our best classifier (Figure 7), while others were chosen by other models (Supp. Fig. 9).

**Figure 7.**
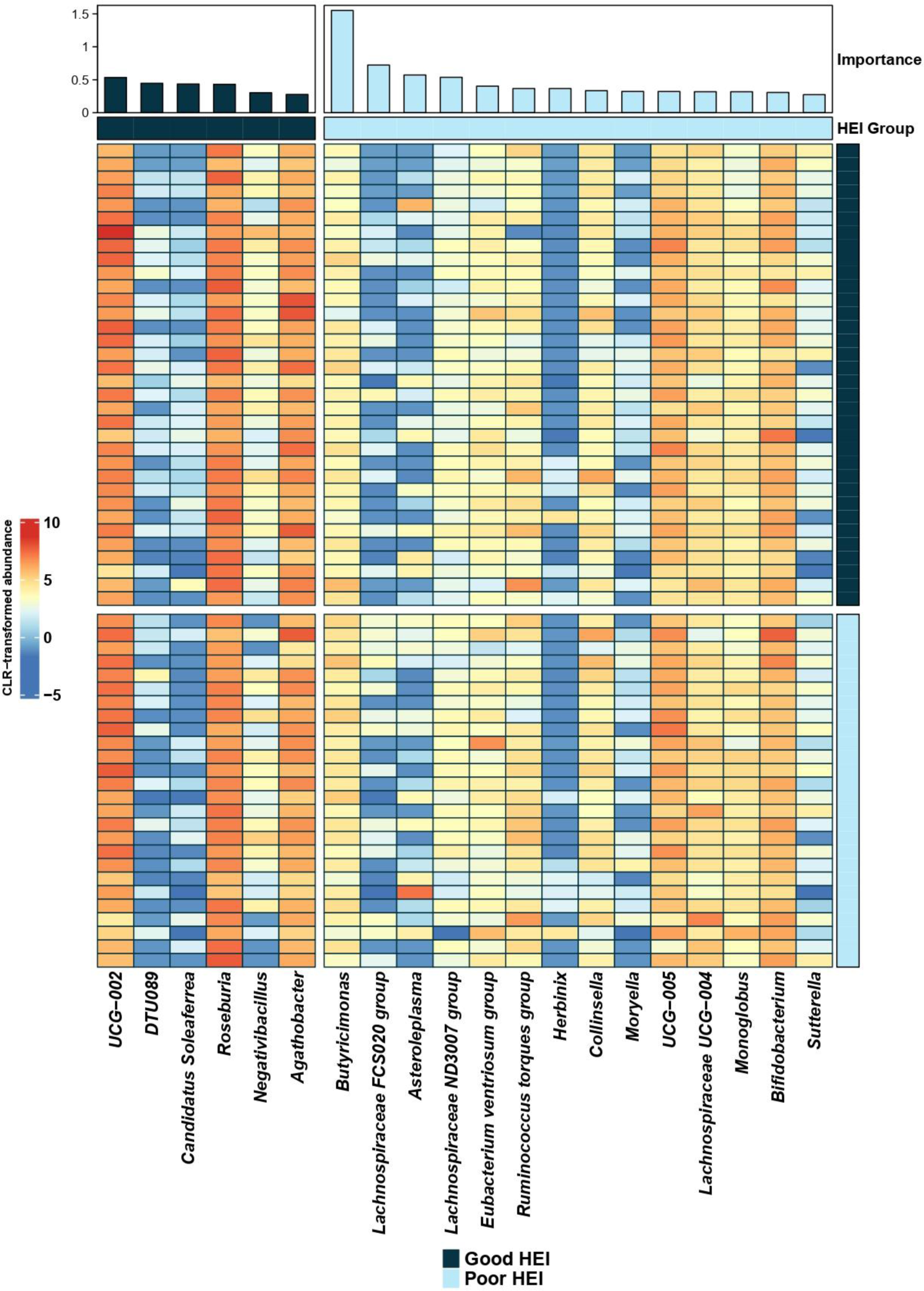
20 most important features according to the RF model based on the HEI classification. Feature importances are plotted at the top of the figure. Bar height indicates feature importance and bar colour indicates whether the genus is enriched in the good HEI group (dark blue) or the poor HEI group (light blue). Taxa are sorted according to the group where they are enriched and in descendent order. A heatmap of CLR-transformed relative abundances is plotted below. The bar that separates both plots and the one on the right indicate the group in which each genus is more abundant and the one the participants belong to, respectively.

Additionally, and given that 7 individuals were found in the study cohort to have T2D family history after reviewing this cohort’s metadata, the aforementioned potential markers were also evaluated in these particular volunteers in order to determine how their abundances varied in comparison to the rest of the participants both for this classification and the previous one. For the HPF classification, 4 of these people belonged to the low consumption group and the rest to the other one; and for the HEI classification 4 people pertained to the good HEI group and the rest to the poor HEI group. Given the division of these individuals between groups in both classifications and abundances following the same tendencies as the rest of the volunteers from this cohort, results are inconclusive for this matter (Supp. Figs. 3 and 10).

Furthermore, the aforesaid taxa, along with the rest, were tested for significant differences in their abundances, using the Kruskal-Wallis test, among participants of our validation cohort clustered in quantiles depending on their HEI and several T2D biochemical markers (HbA1c, HOMA-IR and adiponectin), with the objective of finding potential associations between these genera and these markers used to diagnose T2D and IR. Some differentially abundant taxa among quantiles included *Lachnospira*, *CAG-115* and *Acetatifactor*, all of them highlighted by DESeq2, along with other taxa like *Dialister* or *Pseudoflavonifractor*, which were not identified as important by any of our HEI-based classifiers but had been underlined as such by some of our HPF-based classifiers (Supp. Fig. 2). Moreover, some of these differentially abundant taxa followed expectable tendencies in their abundances along with these markers’ levels (e.g., *Bacteroides*’ abundance tended to increase with lower values of HbA1c (%)), while this did not happen for other genera (Supp. Figs. 12–15).

## Discussion

In this study, GM of 60 participants of a cohort, named “Study cohort”, derived from a study carried out at the IMDEA Computational Biology group, as well as 46 participants (“Validation cohort”) of another study developed at the same group, was analysed. The first one’s volunteers were classified, using these GM data, according to their HPF consumption in the first place, and then depending on their HEI, on the basis of which the participants of the validation cohort were also clustered. HPF consumption and HEI are factors that were paid attention to since, on the one hand, unhealthy diets are characterised, among other aspects, by a higher consumption of industrially processed products, which has been associated with several health issues [5], [20]. On the other hand, some studies have found a clear association between HPF consumption and HEI [26]–[28], as well as between the latter and GM [114], sparking thereby our interest in the link existing among these three.

As for demographic and clinical parameters, no significant differences among individuals were found for none of the classifications and for none of the evaluated cohorts, as well as regarding physical activity, alcohol consumption and smoking. The same results were obtained when testing for differences between the HEI-based groups from the validation cohort as for some T2D biochemical markers. When consulting the literature, there are previous studies in which statistically significant differences are found among HPF consumption groups for these variables [115]–[117]. Nonetheless, they perform these analyses on cohorts with much higher sample sizes than ours. As for HEI, evidence published in the literature is inconclusive, since in some cases HEI has been associated with BMI [118] but there is also proof of BMI not experiencing any significant changes when grouping participants according to their HEI [119], which could evidence thus the difficulty underlying an appropriate choice of cut-offs for the definition of HEI-based groups in studies similar to ours. Regarding physical activity, alcohol consumption, smoking and some T2D biochemical markers’ levels, our results agree with the bibliography [119]. Given these facts, further studies should be performed in the future in order to determine more precisely the links between HPF consumption, HEI and these variables.

Regarding our diversity analyses, no significant differences were found between groups for any of the classifications of both cohorts. These results agree with the ones from previous studies, in which no significant differences are evidenced as for these diversity metrics [5], [120]. However, other studies report opposite results, indicating a negative association between HPF consumption and alpha/beta diversity [121]. This, on the whole, suggests that the link between HPF consumption and GM homeostasis remains unclear [122]. As for HEI, our results might be due to HEI being inappropriate as a variable for the detection of differences between microbial ecosystems as poor diets that are diverse but score similarly may mask trends due to certain dietary constituents [114].

Several binary ML classifiers were also built with the purpose of classifying volunteers from our study cohort into two categories depending on their HPF consumption or their HEI, based on their GM composition. These models were, firstly, trained with centred log-ratio transformed relative abundances, since CLR-transformation converts compositional data to scale-invariant data and captures the relationships between features, normalising thus the data and stabilising the variance [123]–[125]. Secondly, they were generated by splitting randomly the data and using LOOCV, since this method has far less bias in comparison to other CV procedures and causes no randomness in the training/validation set splits [104], apart from several studies using this approach for the classification of metagenomic samples [126]–[128].

In general, given our results, it can be seen that models built using the mikropml R package perform better than those built with caret, and those shown in section 4 had reasonably satisfactory AUCs for their corresponding test sets. However, the considerably wide 95% confidence intervals for all models should be taken into account, since this indicates less precision and a larger margin of error [104], [107]. In light of this, in order to increase the robustness of our models, some strategies that could be assessed in future studies might be the usage of compositional kernels designed specifically for compositional data [129], [130] or deep learning-based classifiers [66], [131], [132] or frameworks [61], [63], [133]–[135].

Once all classifiers for the HEI classification of the study cohort were built, the one created using mikropml and the RF algorithm was used for validation, since it had yielded the highest AUC, of 0.85, over the test set among the rest of the classifiers. Subsequent validation of this model on the validation cohort revealed a decrease in the AUC, which was of 0.7. Even though this does not necessarily represent a suboptimal result, it might indicate low generalisability of our model given the considerable drop in the AUC when the model faces an unprecedented dataset, perhaps owing to the model having been trained with a small training set. It has also been suggested that additional data, such as metabolomic or lifestyle-related information, combined with microbial data, might improve prediction accuracy [136], especially when performing disease prediction, something strongly linked to our case, taking the evidence regarding associations between HEI and obesity into account [118], [137], [138], as well as between HPF consumption and several pathologies.

Various taxa were outlined by the best ML model, regarding the HPF classification, to be the most relevant. This model, along with other 3, highlighted the importance of the *Prevotellaceae NK3B31 group* genus, agreeing in addition to the differential abundance analysis (DAA) performed on DESeq2. This genus has previously been reported to be involved in starch degradation and glucose metabolism, providing energy for growth, reducing inflammation and increasing short-chain fatty acid (SCFA) production, contributing thereby to a balanced gastrointestinal tract [139], [140]. As for the undefined genus linked to the *Muribaculaceae* family, it has been inversely linked to the consumption of foods related to poorer dietary habits, like red meat [141]. Furthermore, several studies in mice have associated this family with body weight loss and a healthier diet after noting an increase in its abundance following a higher consumption of fibre and pulses [142], [143]. Indeed, this family has been shown to degrade complex carbohydrates [144] and decrease its abundance in obese mice [145] while increasing under high fibre intake [146]. Moreover, *Rikenellaceae RC9 gut group*, another genus identified in this work to be differentially more abundant in the group with lower HPF consumption, contributes to the digestion of crude fibre [147], [148] and the protection against oxidative stress [149]. Moreover, these genera’s abundances have also been reported to decrease in subjects with T2D [147]. On the whole, this ML model identified some taxa that could be biomarkers of healthier dietary habits, even though more research is needed in humans.

On the other hand, this ML model also identified other taxa that are more related to a poorer diet, such as *Dorea*. This genus, along with *Collinsella*, has been reported to be more abundant in overweight or obese individuals [150], after evidencing a positive association between this genus, BMI and body weight [151], [152], and has been suggested to contribute to a decrease in the abundance of SCFA-producing bacteria [151], [153]. It has also been linked to colorectal cancer (CRC) [66], a higher consumption of fried meat [151] and prediabetes [154]. *Dialister*, another genus that was identified in this work to be differentially more abundant in the group with a higher HPF consumption, has also been linked to weight gain [155] and T2D [156]. *Phocea* and *Pseudoflavonifractor*, identified by some of our HPF-based models as well, have also been related to T2D [157]. All genera presented in this paragraph belong to the *Firmicutes* phylum.

In relation to the HEI classification, various taxa were outlined by the best ML model to be the most relevant. One of these was *Butyricimonas*, which belongs to the *Bacteroidetes* phylum and was more abundant in the group with poorer HEIs. This genus has been positively associated with the adoption of a high-fat diet [158], but also with normal BMIs in another study [159]. Moreover, the *B. virosa* species has previously been stated to have a beneficial effect on obesity and T2D by promoting SCFA production [160] and thus the secretion of the glucagon-like peptide-1 (GLP-1) [161], an incretin that activates glucose-dependent insulin secretion and inhibits glucagon release [162]. Conversely, *Sutterella*, which was more abundant in the group with poorer HEIs, has been reported to be positively correlated with the likeness of developing T2D [163]. *Lachnospiraceae FCS020 group*, despite being more abundant in the same HEI group, has been associated with protective properties and a healthier diet in mice [141]. *Roseburia*, which was more abundant in the group with healthier HEIs, has been widely reported to decrease its abundance in T2D patients [164].

Furthermore, *Gastranaerophilales* and *Intestinimonas*, identified to be differentially abundant in the group with healthier HEIs from the study cohort, have been associated with anti-inflammatory effects [165] and a decrease in abundance in prediabetic patients [166], respectively. Conversely, *Erysipelotrichaceae UCG-003* was more abundant in the group with lower HEIs and has indeed been linked to CRC and inflammation [44], while *Akkermansia*, indicated by DESeq2 to be differentially abundant in the group with higher HEIs from the validation cohort, has been inversely associated with obesity and T2D [167]. In summary, even though some of our results, regarding both our ML models and DAA, agree with the literature, further research should be performed to obtain more robust conclusions, especially in the case of taxa that have been evaluated in studies with animal models but not in humans.

Several taxa that could be markers of HPF consumption and, in general, poorer dietary habits, were therefore identified. Moreover, most of these genera are associated, one way or another, with T2D. In fact, GM and alterations in its composition have been associated in multiple studies with the pathogenesis of T2D, for example after evidencing decreased abundances of butyrate-producing bacteria and increased opportunistic pathogens in patients with T2D [168]. A promotion of metabolic endotoxemia, insulin resistance and hyperglycaemia has also been reported in T2D patients as cause of gut microbial dysbiosis [169]. GM has an influence on the response for some antidiabetic agents as well, such as metformin [170] or GLP-1R agonists (GLP-1RAs) [170], contributing to heterogeneous responses to these treatments. GLP-1RAs, for instance, have proven to alter GM composition in mice [171] and humans [172], increasing the abundance of taxa with immunomodulation effects like *Roseburia* or *Bacteroides* and decreasing that of genera with pro-inflammatory properties such as *Dialister* [172].

Even if there were only 7 individuals in our study cohort with T2D family history and no information on this matter in the validation cohort, the aforementioned alterations on T2D host metabolism might explain why some of these participants, despite being young (considering T2D affects mostly adults aged 45-64 [173]) and having normal BMIs and good eating habits, still had lower abundances of certain taxa inversely linked to T2D, such as *Prevotellaceae NK3B31 group*, among others, as seen in Supp. Figs. 3 and 10. GM might thus have a negative influence on these volunteers’ T2D susceptibility, despite them following a health-conscious lifestyle, due to this family history. In fact, it would be of interest in future studies to evaluate some of our markers in a bigger cohort of individuals with T2D family history, in order to find out whether people with this history, despite having wholesome habits, could still present higher abundances of T2D-related taxa, suggesting a predisposition for T2D induced by GM.

On the other hand, several T2D-related biomarker data were collected from the validation cohort, which were used to group them in quantiles. All features identified in their GM data were tested for significant differences in abundance using the Kruskal-Wallis test and some significant taxa were then plotted, as shown in Supp. Figs. 12–15. Even if some expectable tendencies can be seen, like the increase in *Pseudoflavonifractor*’s abundance along with HOMA-IR or the decrease in *Bacteroides*’ abundance with increasing HbA1c (%) levels, *Dialister*’s abundance did not increase, for example, along with HbA1c (%) values. Given these inconclusive results, these taxa should be further studied to discover likely associations of GM with these biochemical markers or confirm their link with T2D.

The main limitation of this work is the small sample size for both cohorts, since this has a direct effect on all microbiome analyses and the ML models’ robustness. Furthermore, the project from where our study cohort derived was cross-sectional and, as for the study from which our validation cohort came, even though GM composition data were collected at three time points, only data from the first visit were taken into account for our work because of space and time constraints. It would have been interesting to assess changes in GM composition over time for the same cohort, instead of comparing microbial biomarker candidates, identified using ML models trained on data coming from another study, between two different cohorts. This would have also allowed us to confirm with more conviction whether our biomarkers are indeed associated with diet, more specifically with HPF consumption, and diabetes. Additionally, cut-offs used to cluster participants from both cohorts in two groups, for both classifications, were chosen mainly with the purpose of getting as much balance as possible as for the groups’ size. Other classification strategies should be assessed in order to seek more significant results.

Regarding the ML classifiers that were built, some additional strategies could be performed in future work to increase the robustness of the candidates that were identified as markers in this work. Even though additional feature selection techniques are discouraged when using the caret R package, several studies have implemented them using other R packages or the scikit-learn python package [174]. In addition, even if four different ML algorithms were tested and, to the best of our knowledge, the caret and mikropml R packages were compared for the first time, the assemblage of more appropriate or powerful techniques, using deep learning, may yield better results. On the other hand, regarding the DAA that was performed using DESeq2, even if most of the taxa identified by this tool concurred with the literature, there are more available approaches, like edgeR [175], ALDEx2 [176] or ANCOM [177]. A comparison between the results from each one of these techniques might have been convenient in order to select differentially abundant taxa determinedly; something that in fact will be carried out in a review, still under construction, focused on the application of machine learning to the analysis of microbiome data, in which this work’s author is currently participating as the main author, along with other ML4Microbiome COST Action (CA18131) members.

## Conclusions

In this work, we aimed to improve our understanding of how diet and, specifically, HPF consumption affect human health by studying changes in GM composition. By carrying out DAA and building several ML models, different genera that could be potential biomarkers of healthy/unhealthy dietary habits, HPF intake and diabetes were identified. Nevertheless, further research, by evaluating GM composition at different time points, using additional ML techniques and recruiting more participants is needed to confirm these observations.

## Declarations

### Ethics approval and consent to participate

Not applicable.

### Consent for publication

Not applicable.

### Competing interests

The authors declare that they have no competing interests.

## Funding

This work has been carried out under the context of the projects AI4FOOD (Y2020/TCS-6654) Artificial Intelligence for the Prevention of Chronic Diseases through Personalized Nutrition financed by the 2020 call for Synergic R&D projects, of the Community of Madrid, and a Proof of Concept Grant from the European Institute of Innovation and Technology (PoC-47). L.J.M.-Z. is supported by Juan de la Cierva Grant (IJC2019-042188-I) from the Spanish State Research Agency of the Spanish Ministerio de Ciencia e Innovación y Ministerio de Universidades. Part of this publication is based upon work from COST Action COST CA18131/ML4Microbiome supported by COST (European Cooperation in Science and Technology).

## Author’s contributions

Conceptualisation, L.J.M.-Z. and E.C.d.S.P.; methodology, V.M.L.M., B.L.-P. and L.J.M.-Z.; writing—original draft preparation, V.M.L.M. and L.J.M.-Z.; writing— critical review and editing, L.J.M.-Z. and E.C.d.S.P.; funding acquisition, L.J.M.-Z. and E.C.d.S.P. All authors have read and agreed to the published version of the manuscript.

## Corresponding authors

Correspondence to Enrique Carrillo de Santa Pau and Laura Judith Marcos-Zambrano.

## Acknowledgements

The authors wish to extend our thanks to Lidia Daimiel Ruiz (IMDEA Food Institute, Spain), Celia Martínez Pérez (IMDEA Food Institute, Spain) and all Computational Biology group members for their scientific comments and fruitful discussions during the development of this work.

## List of abbreviations

16S rRNA: 16S ribosomal RN
ASV: amplicon sequence variant
AUC: area under the curve
BMI: body mass index
CLR: centred log-ratio
CVD: cardiovascular disease
DAA: differential abundance analysis
DALY: disability adjusted life year
DGA: dietary guidelines for Americans
FDR: false discovery rate
GB: gradient boosting
GLM: generalised linear model
GLP-1: glucagon-like peptide-1
GLP-1R: glucagon-like peptide-1 receptor
GM: gut microbiome
GPAQ: global physical activity questionnaire
GTDB: genome taxonomy database
HEI: healthy eating index
HPF: highly processed food
IBD: inflammatory bowel disease
log2FC: log2 fold change
MET: metabolic equivalent
ML: machine learning
NCD: non-communicable disease
PCoA: principal coordinate analysis
PERMANOVA: permutational multivariate analysis of variance
RF: random forest
ROC: receiver operating characteristic
SCFA: short-chain fatty acid
SGB: species-level genome bins
SSB: sugar-sweetened beverage
SVC: support vector classifier
T2D: type 2 diabetes
WGS: whole-genome shotgun sequencing

## Supplementary Material

**Supplementary Figure 1.**
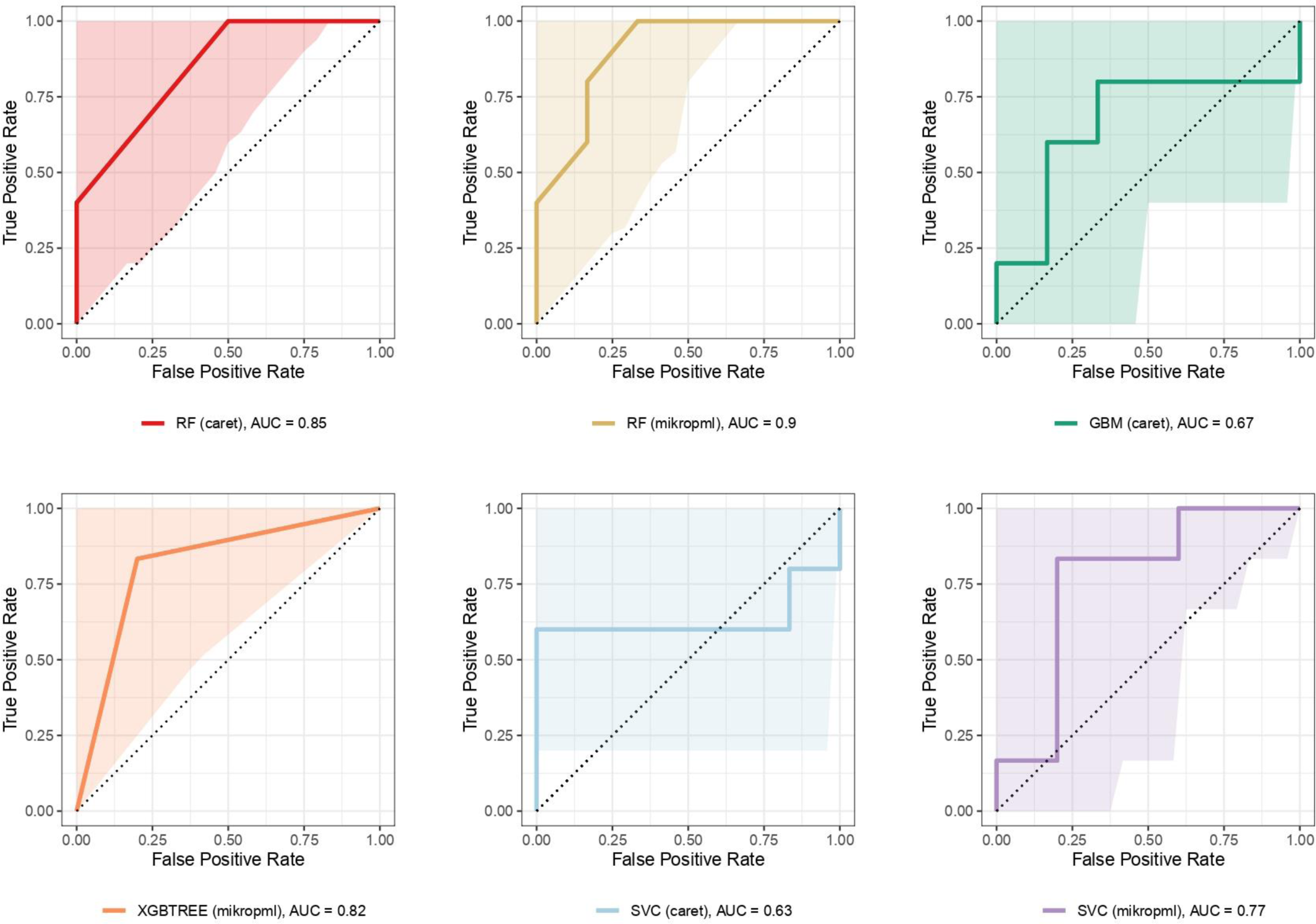
ROC curves for all the binary ML models trained on the CLR-transformed relative abundances table from the study cohort classified according to their HPF consumption. These models were obtained with the hyperparameter values shown in Supp. Table 1. Shadowed areas represent 95% confidence intervals for sensitivity.

**Supplementary Figure 2.**
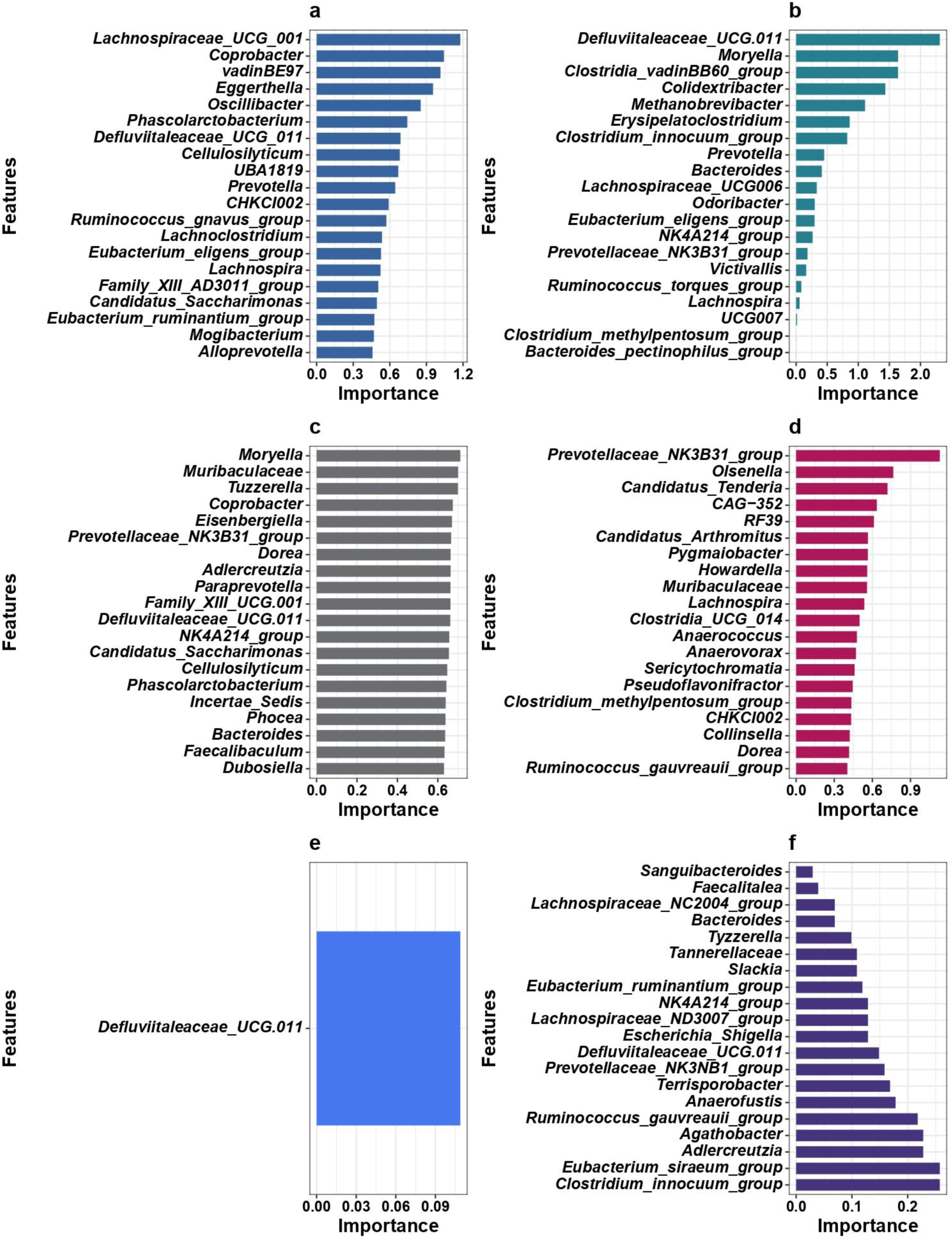
Top 20 features by importance in all binary ML models trained on the CLR-transformed relative abundances table from the HPF-based classified study cohort. **a)** RF (caret) **b)** GBM (caret) **c)** SVC (caret) **d)** RF (mikropml) **e)** XGBTREE (mikropml) **f)** SVC (mikropml). Importances are measured using the mean decrease in the Gini coefficient, except for **c)**, for which feature importances are measured using the ROC curve for each predictor, and **e)** and **f)**, for which they are measured by means of *p*-values, which reflect the probability of obtaining the actual performance value under the null hypothesis (the feature is not important for model performance).

**Supplementary Figure 3.**
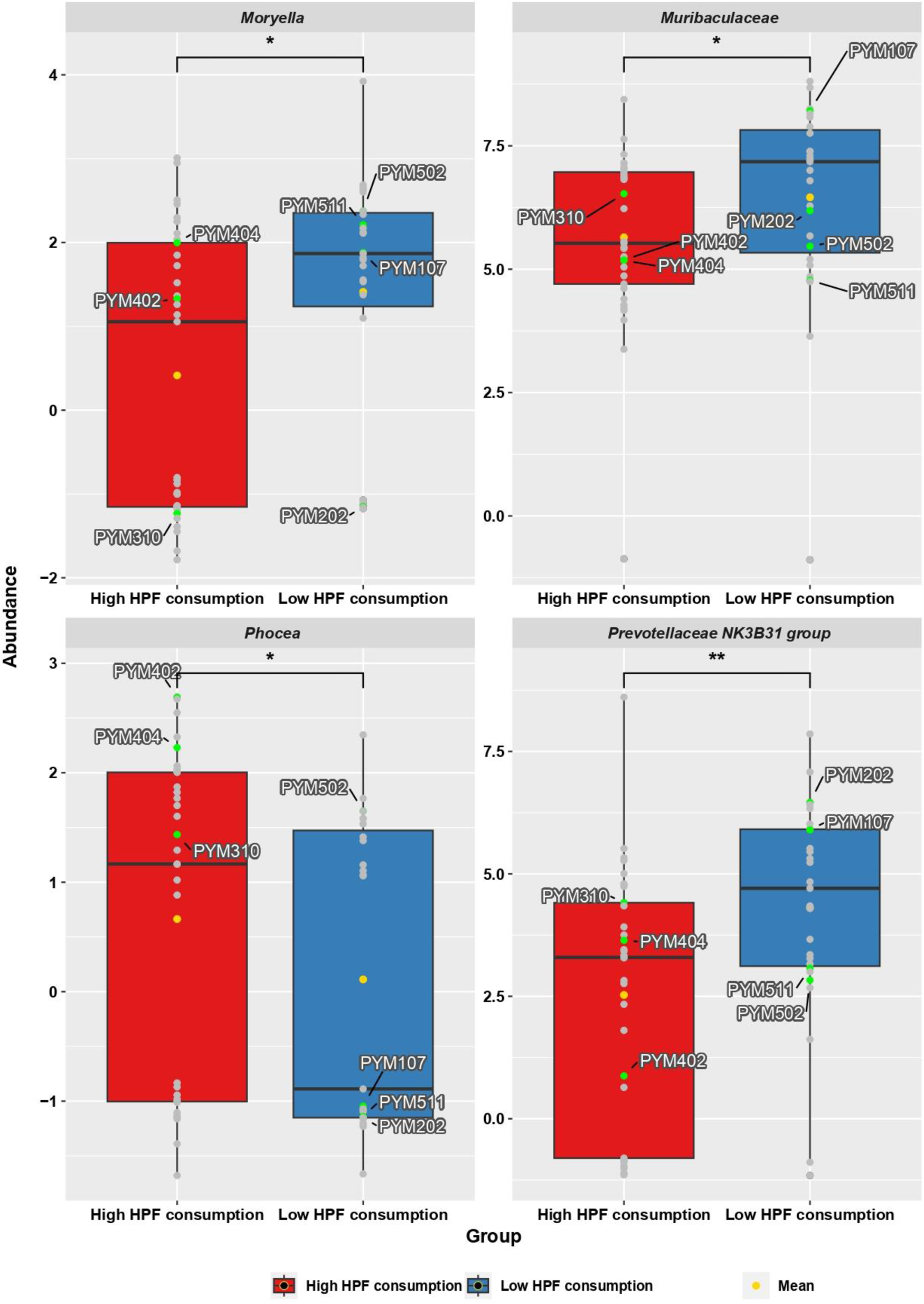
Genera significantly different in abundance between HPF consumption groups (study cohort), according to the Wilcoxon rank sum test (*Moryella*: *p* = 0.015; *Muribaculaceae* (undefined genus): *p* = 0.015; *Phocea*: *p* = 0.04; *Prevotellaceae NK3B31 group*: *p* = 0.009). Mean is indicated in yellow in each group. Individuals with T2D family history are shown in green along with their sample IDs, while the rest of participants are shown in grey.

**Supplementary Figure 4.**
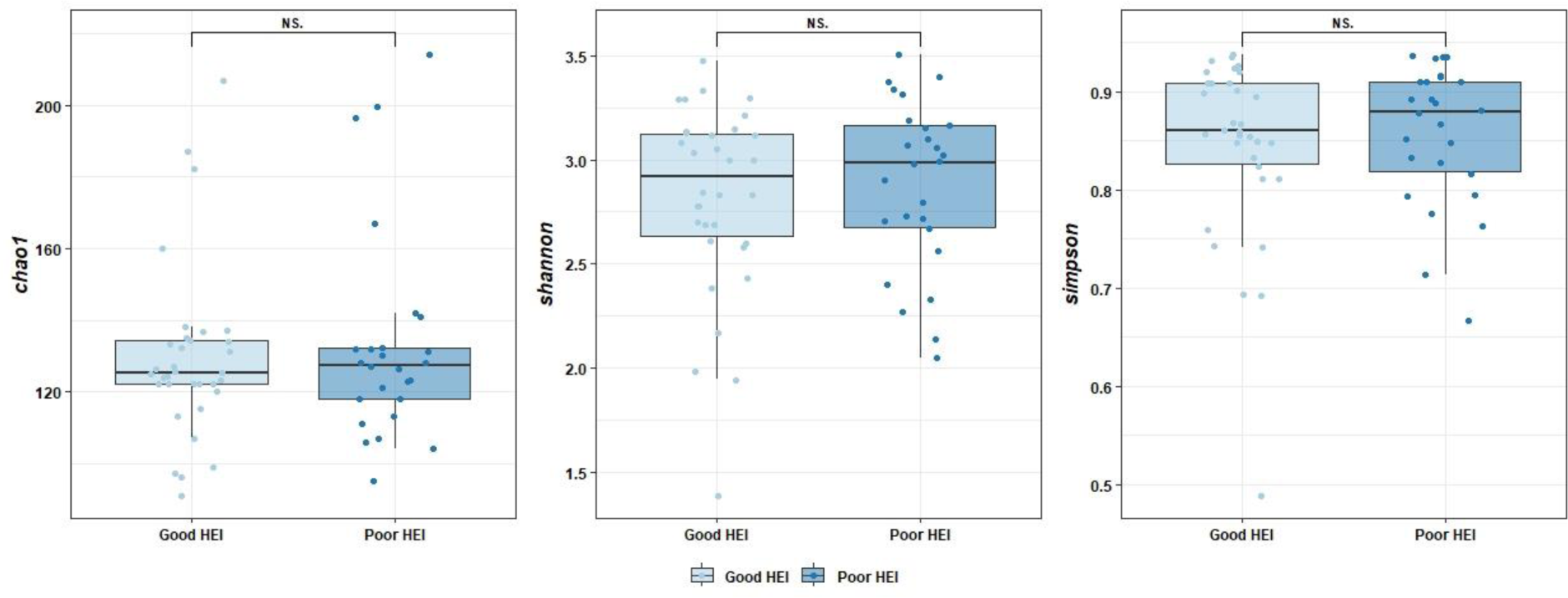
Alpha diversity differences between HEI groups in the study cohort. No significant differences were found in the Wilcoxon rank sum test for the Shannon and Simpson diversity indices (*p* = 0.888 and *p* = 0.888 respectively), as well as the Chao1 richness index (*p* = 0.888). NS.: non-significant.

**Supplementary Figure 5.**
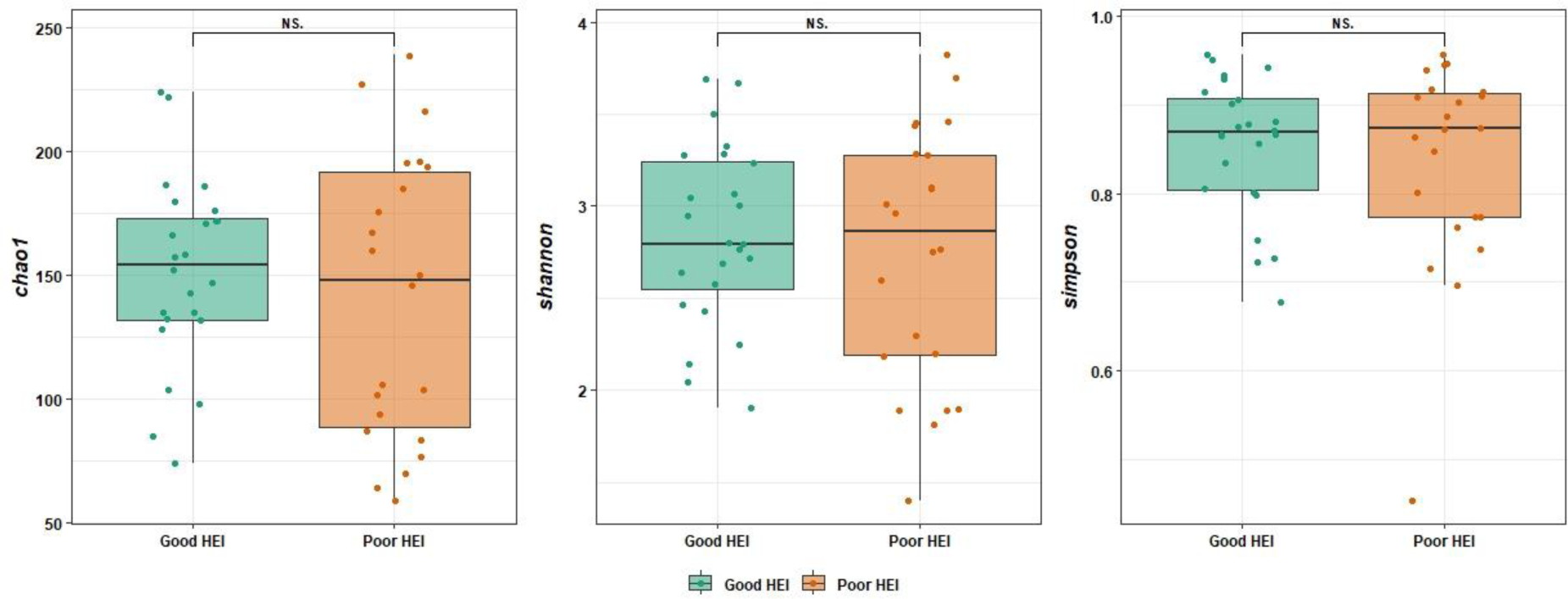
Alpha diversity differences between HEI groups in the validation cohort. No significant differences were found in the Wilcoxon rank sum test for the Shannon and Simpson diversity indices (*p* = 0.957 and *p* = 0.957 respectively), as well as the Chao1 richness index (*p* = 0.957). NS.: non-significant.

**Supplementary Figure 6.**
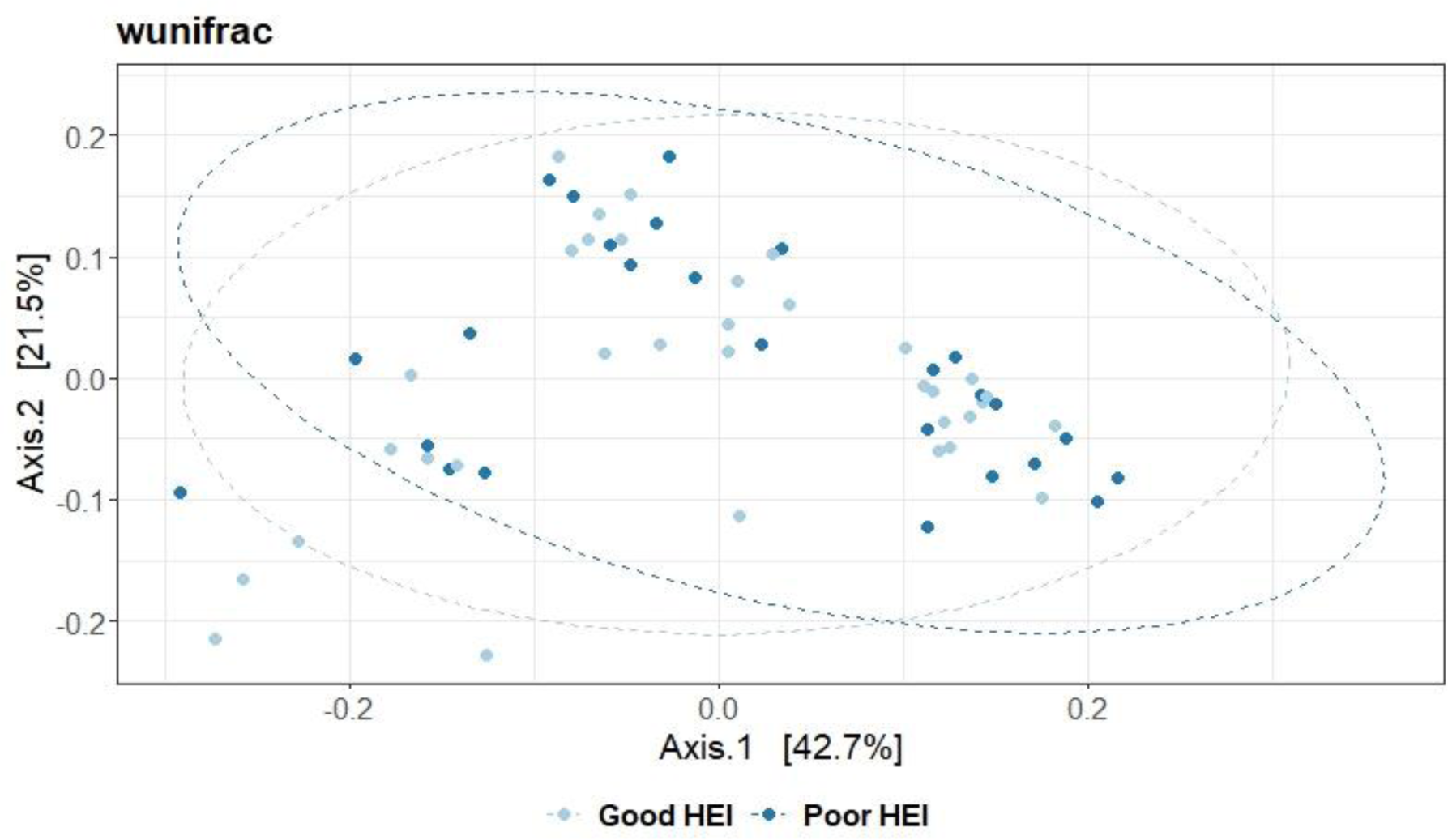
Beta diversity analysis for the study cohort classified based on their HEI: PCoA plot. Weighted UniFrac measurement (*wunifrac*) was calculated with phyloseq and represented via PCoA. Differences between groups were tested with PERMANOVA with 999 permutations. Ellipses represent 95% confidence intervals.

**Supplementary Figure 7.**
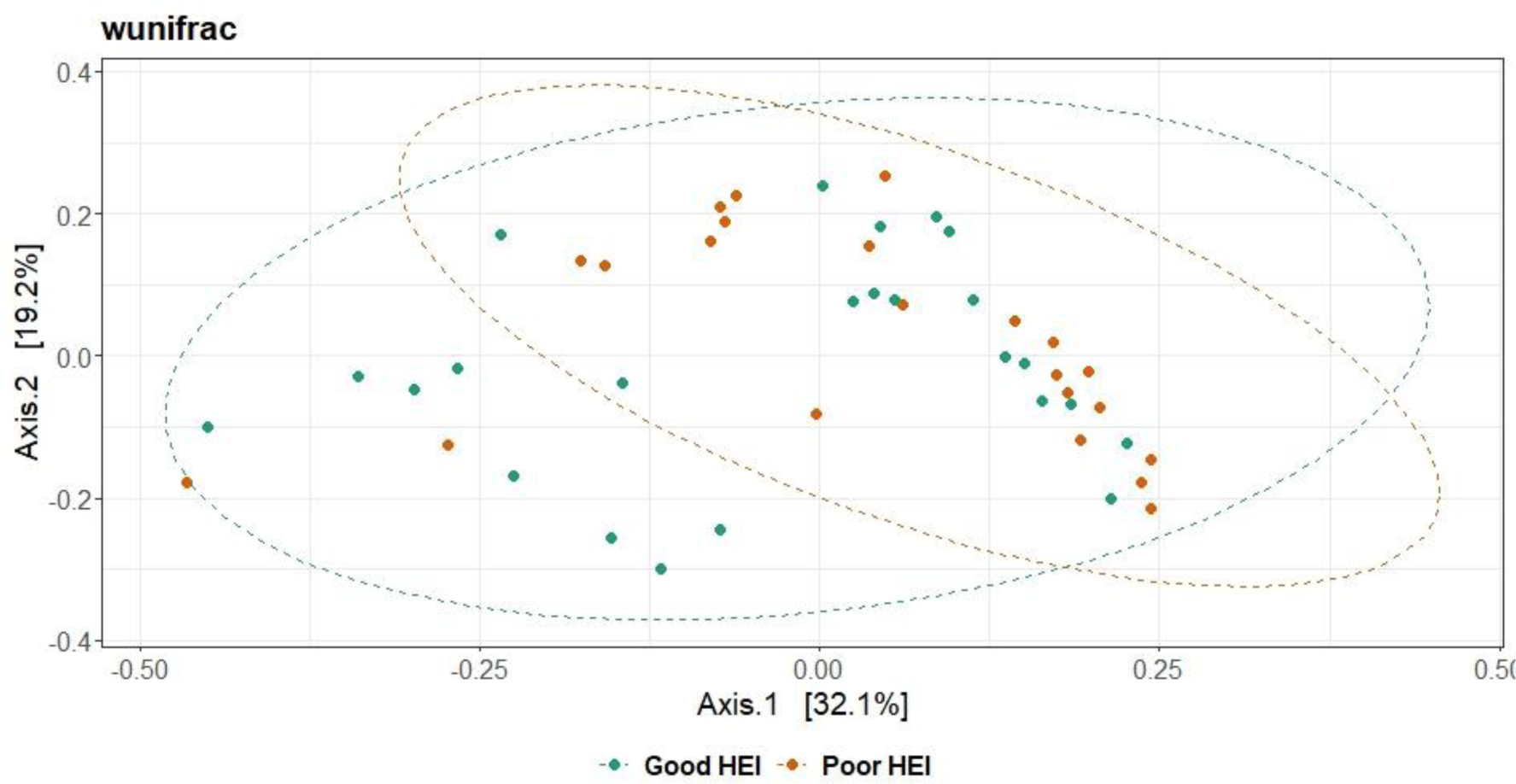
Beta diversity analysis for the validation cohort classified based on their HEI: PCoA plot. Weighted UniFrac measurement (*wunifrac*) was calculated with phyloseq and represented via PCoA. Differences between groups were tested with PERMANOVA with 999 permutations. Ellipses represent 95% confidence intervals.

**Supplementary Figure 8.**
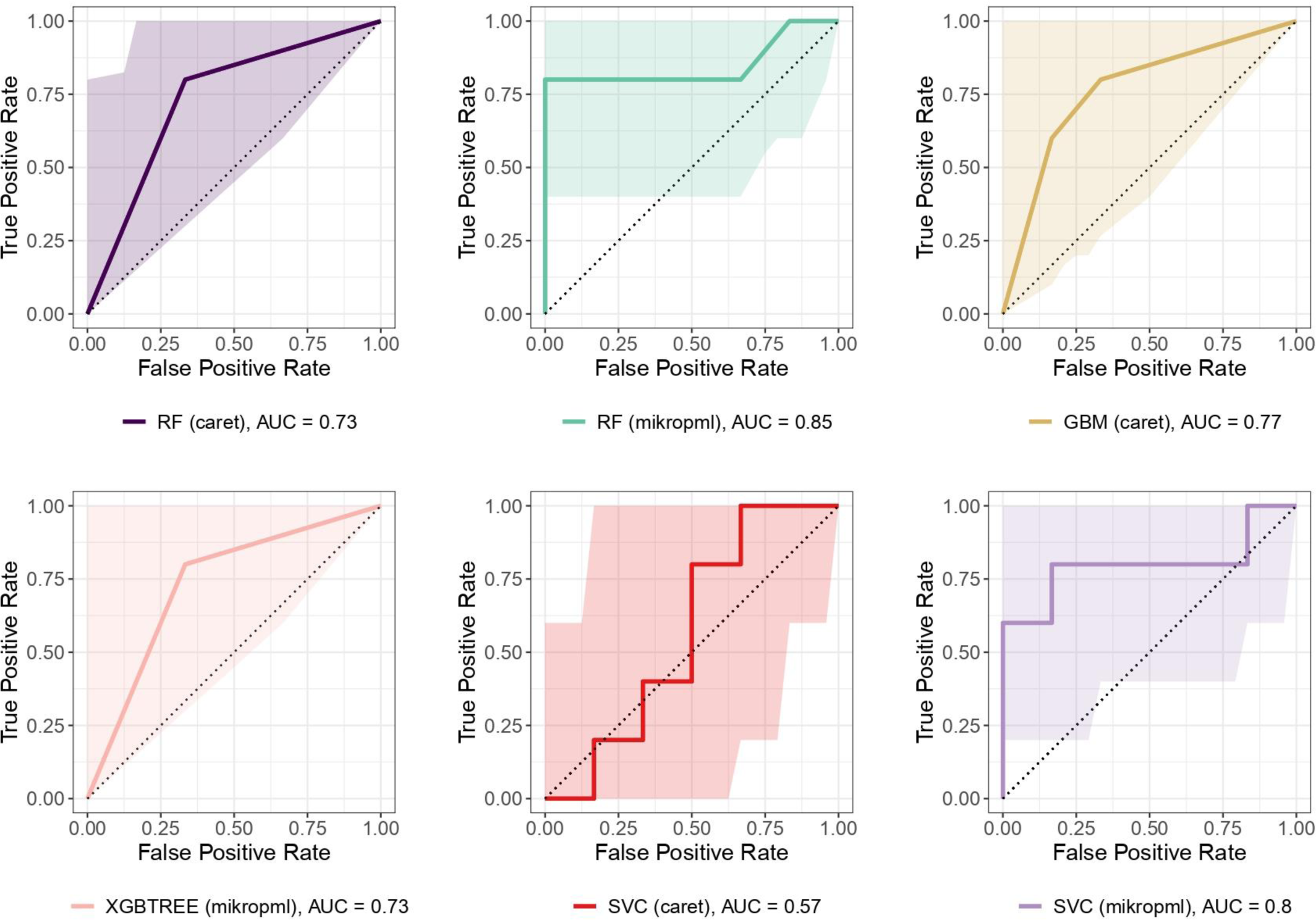
ROC curves for all the binary ML models trained on the CLR-transformed relative abundances table from the study cohort classified according to their HEI. These models were obtained with the hyperparameter values shown in Supp. Table 3. Shadowed areas represent 95% confidence intervals for sensitivity.

**Supplementary Figure 9.**
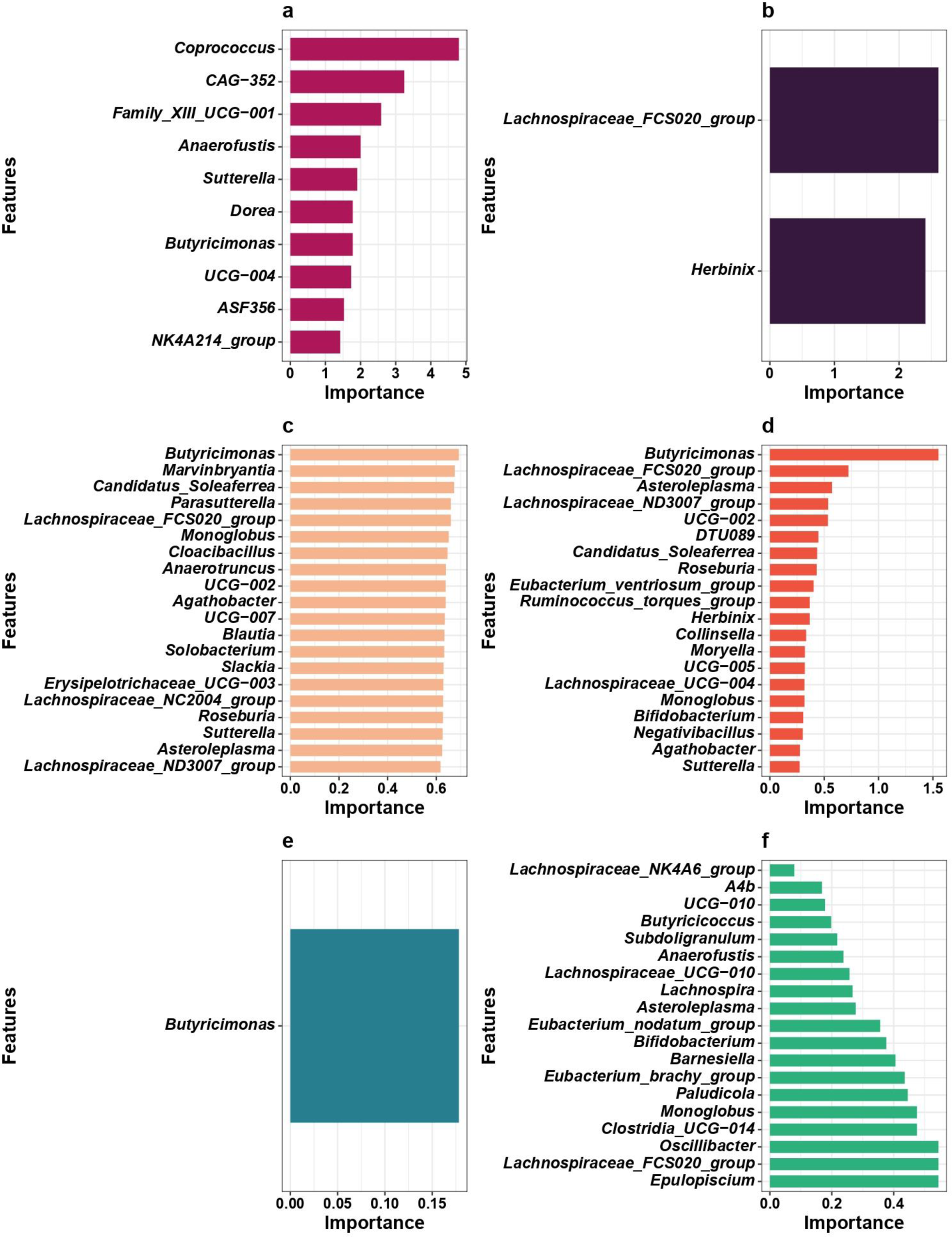
Top 20 features by importance in all binary ML models trained on the CLR-transformed relative abundances table from HEI-based classified study cohort. **a)** RF (caret) **b)** GBM (caret) **c)** SVC (caret) **d)** RF (mikropml) **e)** XGBTREE (mikropml) **f)** SVC (mikropml). Importances are measured using the mean decrease in the Gini coefficient, except for **c)**, for which feature importances are measured using the ROC curve for each predictor, and **e)** and **f)**, for which they are measured by means of *p*-values, which reflect the probability of obtaining the actual performance value under the null hypothesis (the feature is not important for model performance).

**Supplementary Figure 10.**
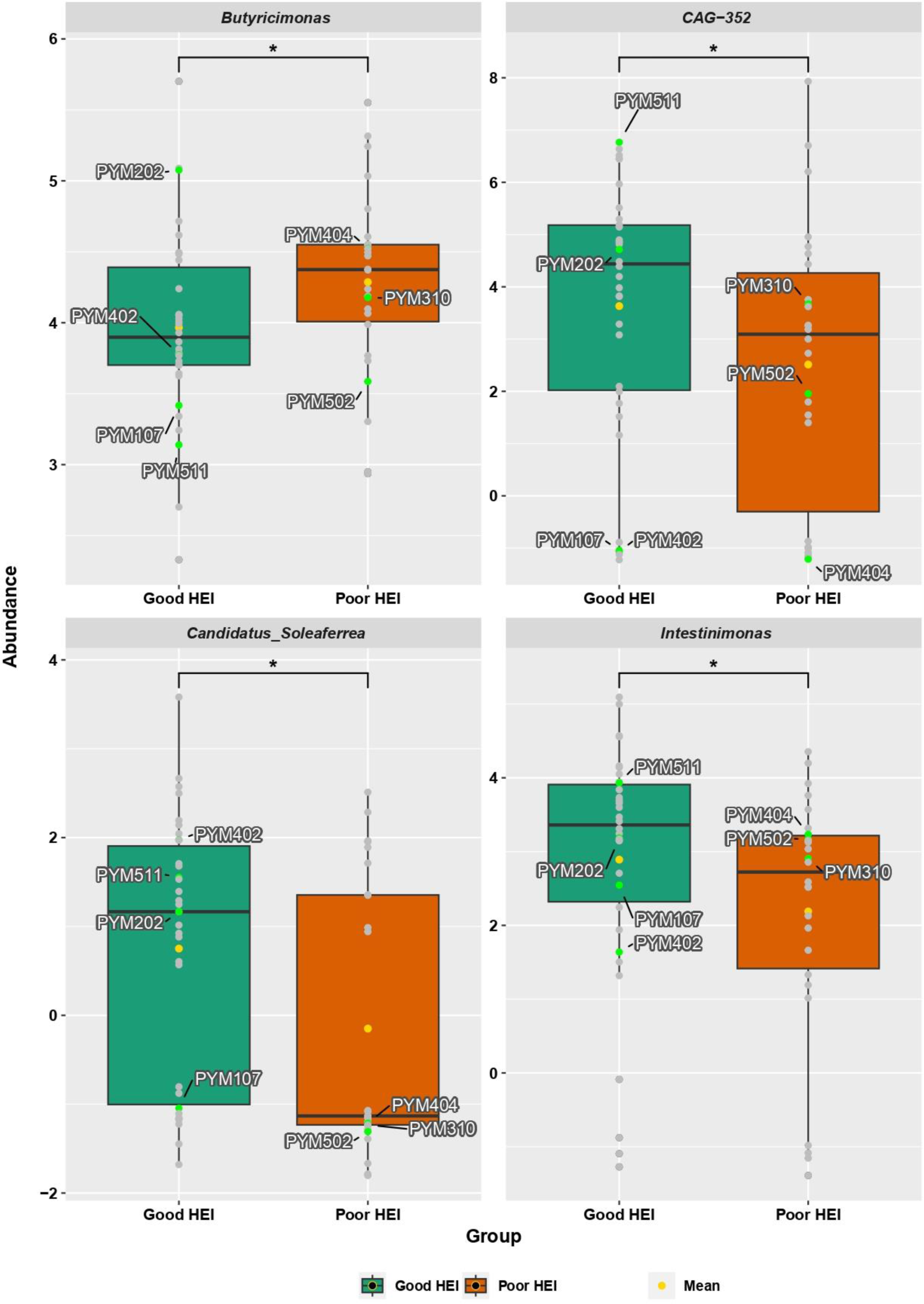
Genera significantly different in abundance between HEI groups in the study cohort, according to the Wilcoxon rank sum test (*Butyricimonas*: *p* = 0.033; *CAG-352*: *p* = 0.048; *Candidatus Soleaferrea*: *p* = 0.023; *Intestinimonas*: *p* = 0.033). Mean is indicated in yellow in each group. Individuals with T2D family history are shown in green along with their sample IDs, while the rest of participants are shown in grey.

**Supplementary Figure 11.**
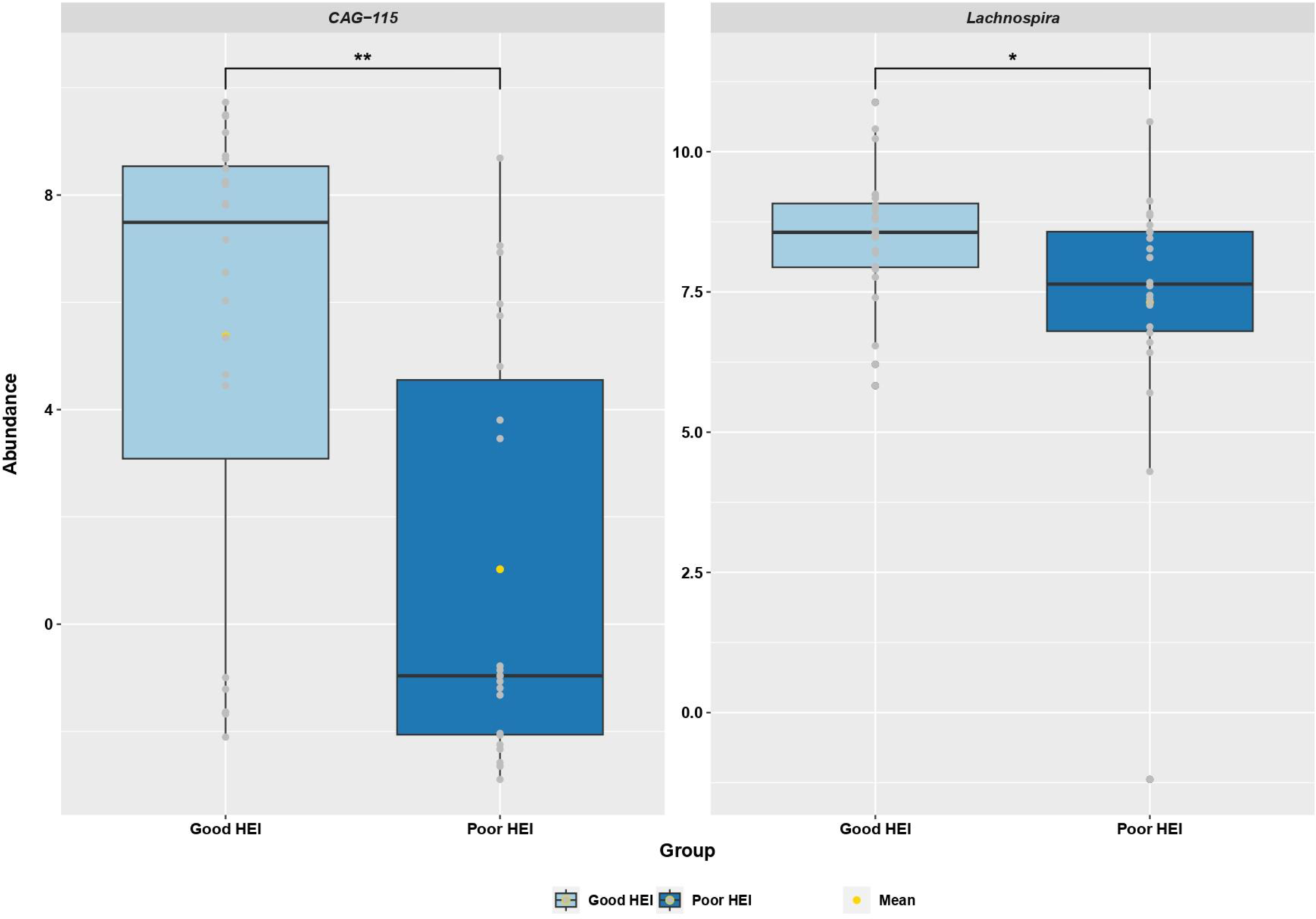
Genera significantly different in abundance between HEI groups in the validation cohort, according to the Wilcoxon rank sum test (*CAG-115*: *p* = 0.001; *Lachnospira*: *p* = 0.028). Mean is indicated in yellow in each group. All participants are shown in grey.

**Supplementary Figure 12.**
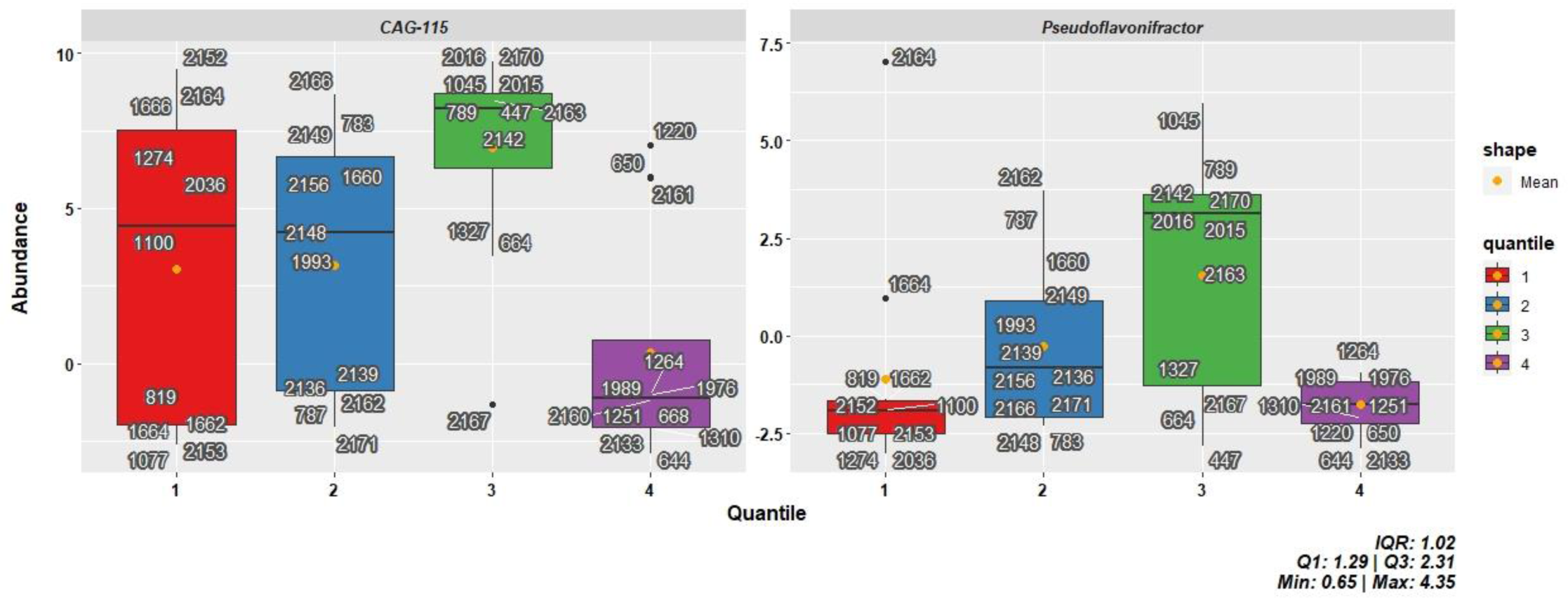
Genera significantly different in abundance among quantiles from the validation cohort obtained based on their HOMA-IR values, according to the Kruskal-Wallis test (*CAG-115*: *p* = 0.0079; *Pseudoflavonifractor*: *p* = 0.0332). Mean is indicated in orange in each group. Sample IDs are shown.

**Supplementary Figure 13.**
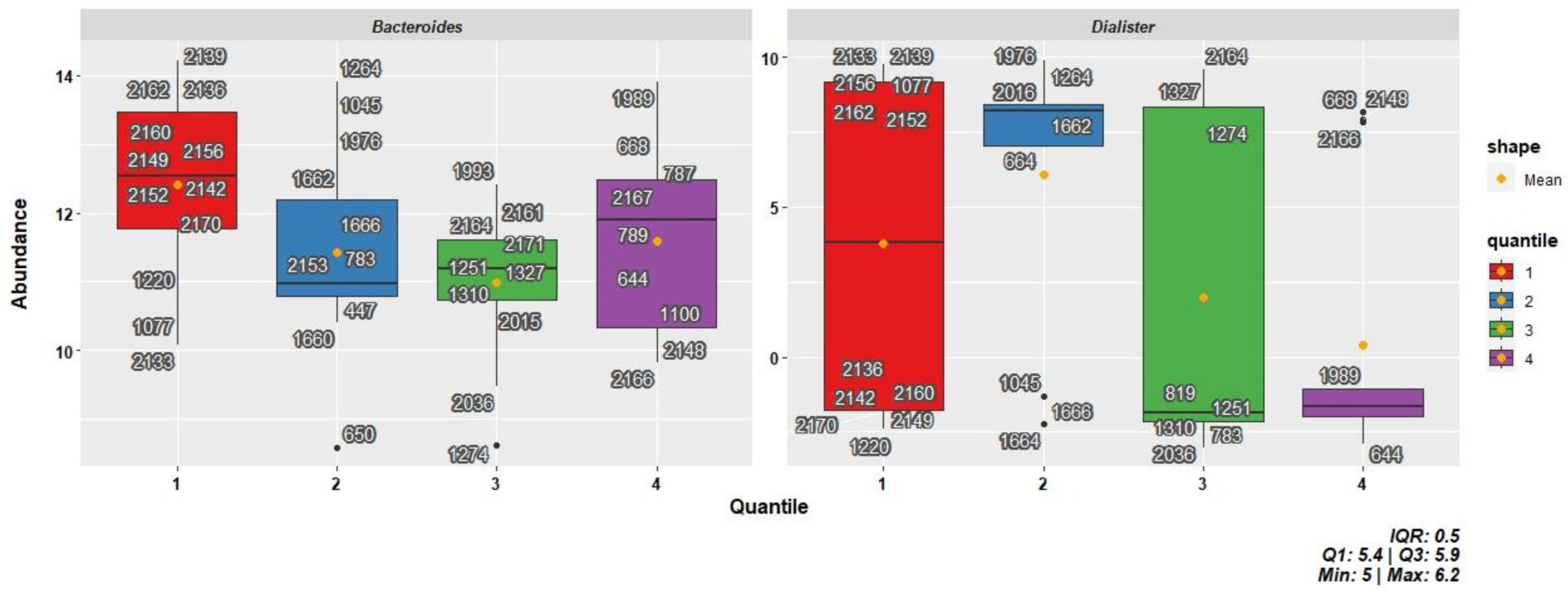
Genera significantly different in abundance among quantiles from the validation cohort obtained based on their HbA1c (%) values, according to the Kruskal-Wallis test (*Bacteroides*: *p* = 0.0486; *Dialister*: *p* = 0.0486). Mean is indicated in orange in each group. Sample IDs are shown.

**Supplementary Figure 14.**
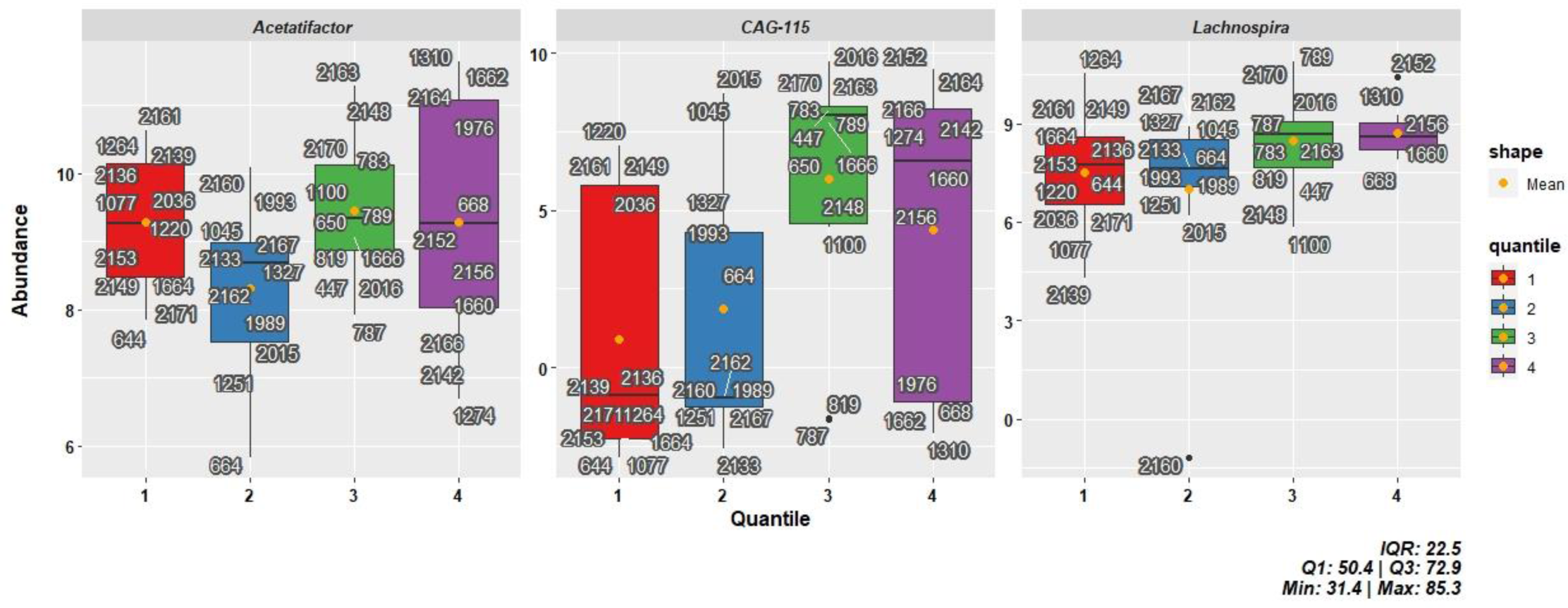
Genera significantly different in abundance among quantiles from the validation cohort obtained based on their HEI values, according to the Kruskal-Wallis test (*Acetatifactor*: *p* = 0.0182; *Lachnospira*: *p* = 0.0348; *CAG-115*: *p =* 0.0328). Mean is indicated in orange in each group. Sample IDs are shown.

**Supplementary Figure 15.**
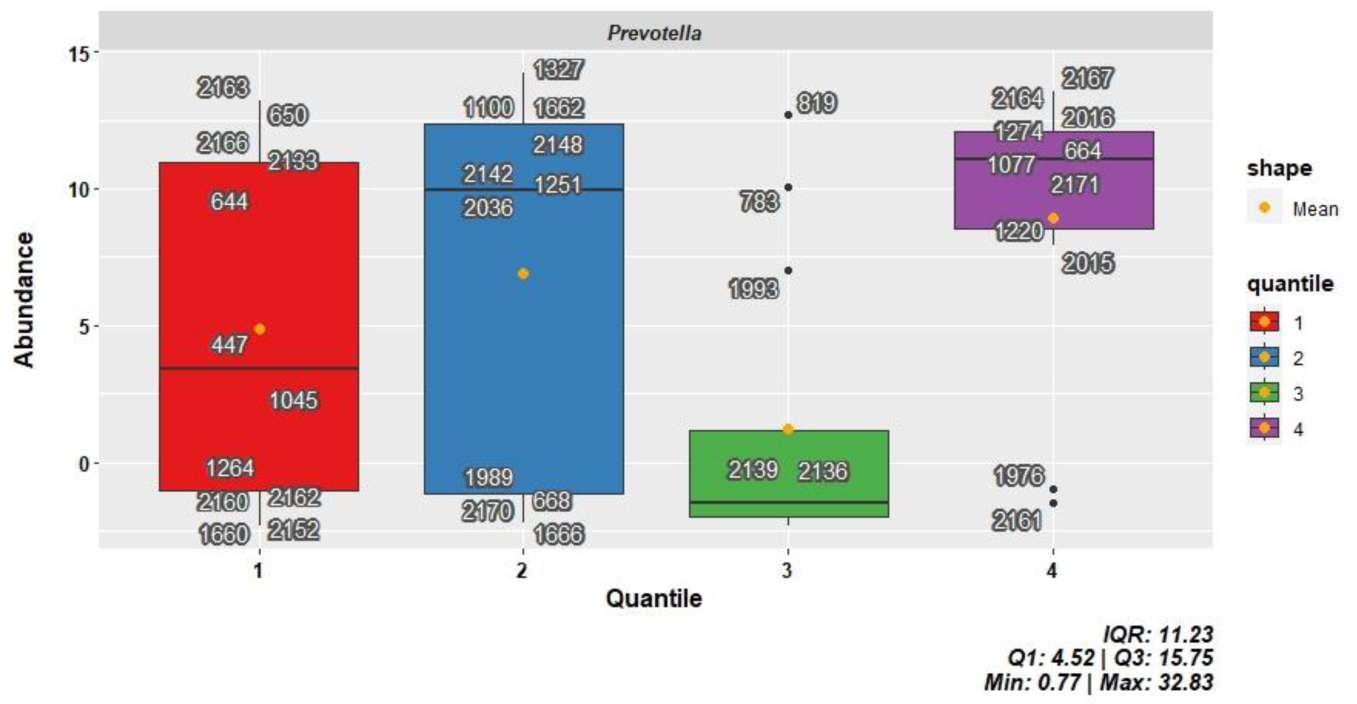
Significantly different in abundance genus among quantiles from the validation cohort obtained based on their adiponectin levels, according to the Kruskal-Wallis test (*Prevotella*: *p* = 0.0414). Mean is indicated in orange in each group. Sample IDs are shown.

**Supplementary Table 1.**
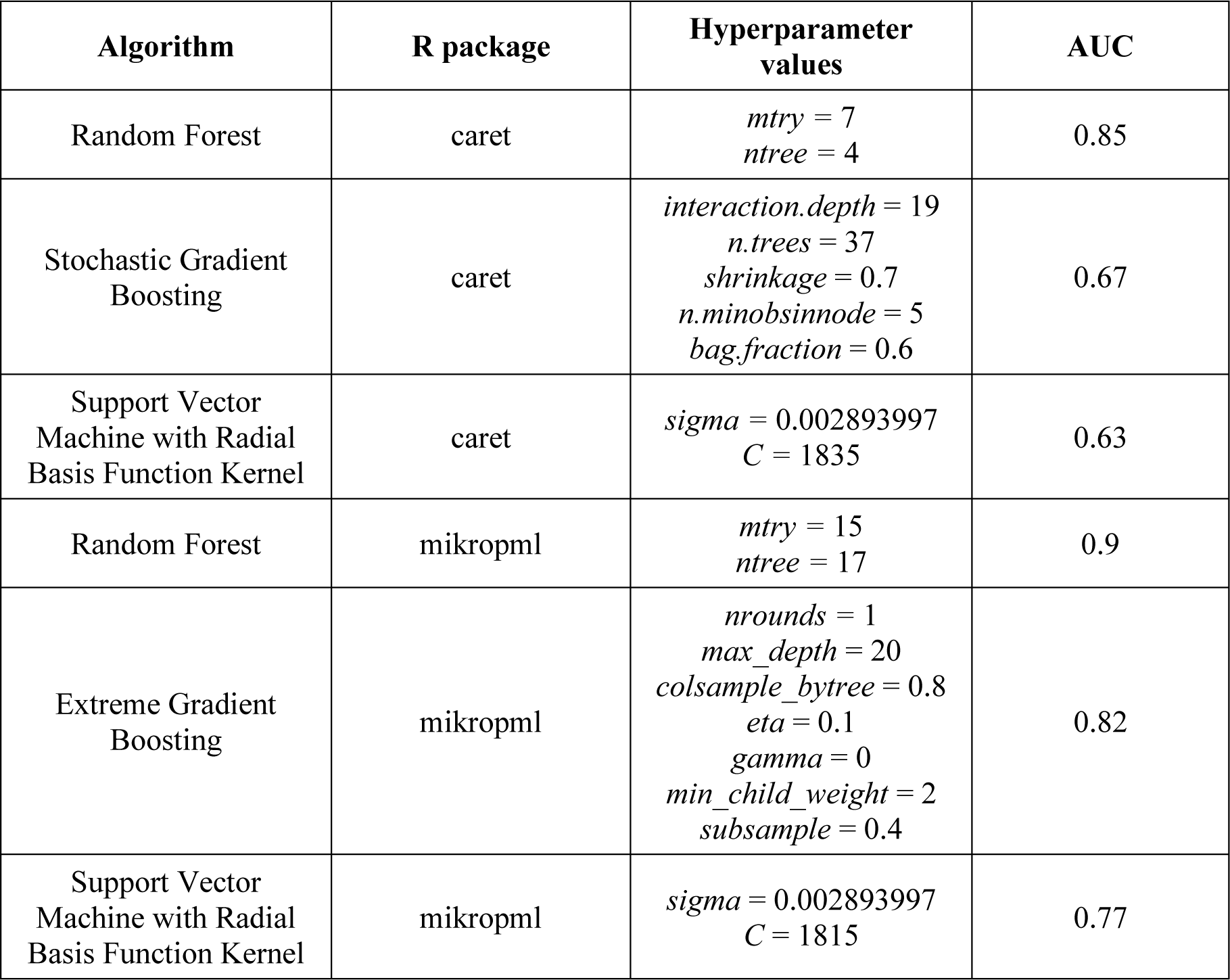
Binary ML models trained on the CLR-transformed relative abundances table from the study cohort classified according to their HPF consumption. Hyperparameter values used to obtain the most precise model for each algorithm and R package are shown, along with their corresponding AUC.

**Supplementary Table 2.**
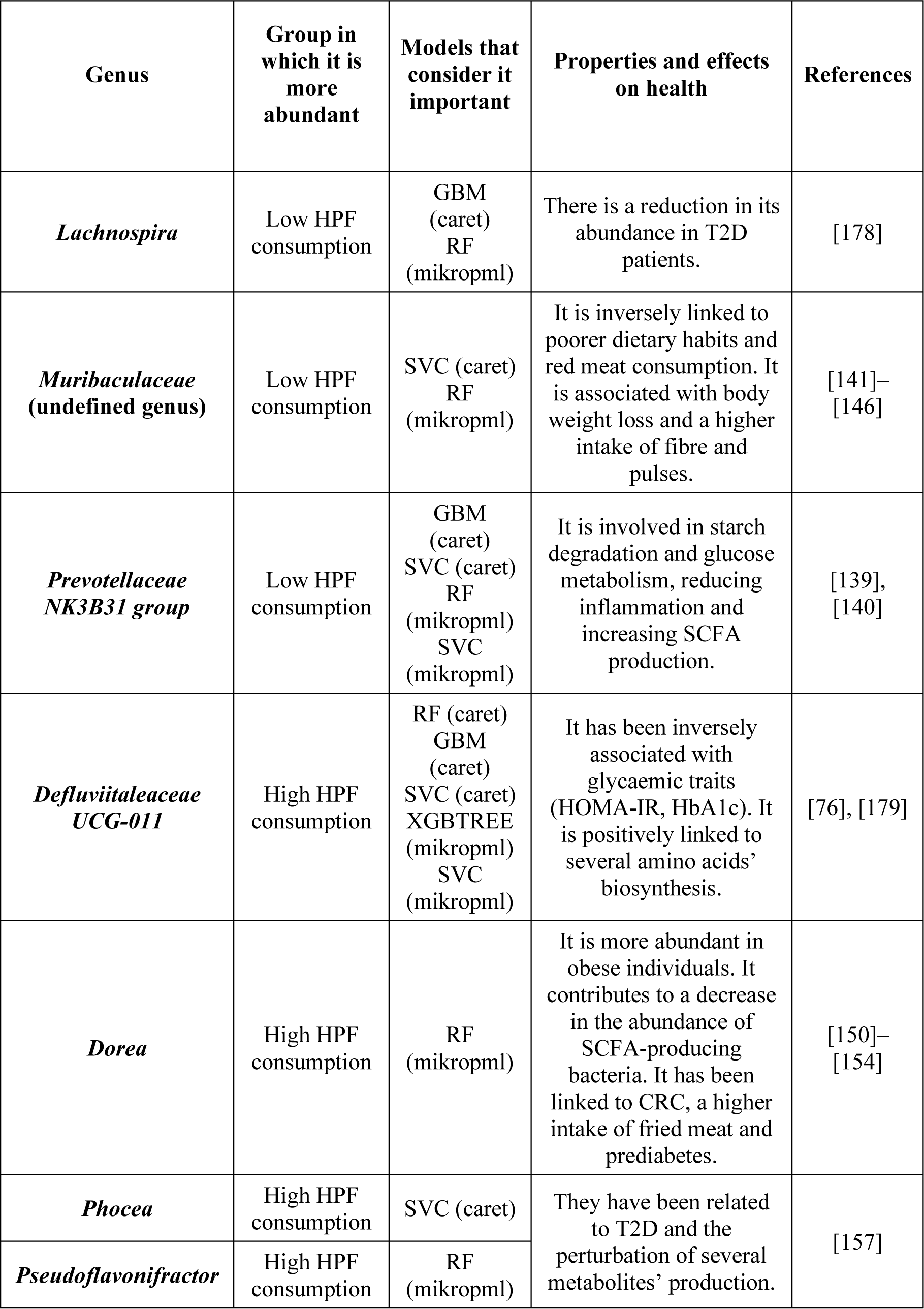
Genera considered to be important by some ML classifiers trained on CLR-transformed GM data from the study cohort, classified based on their HPF consumption.

**Supplementary Table 3.** Binary ML models trained on the CLR-transformed relative abundances table from the study cohort classified according to their HEI. Hyperparameter values used to obtain the most precise model for each algorithm and R package are shown, along with their corresponding AUC.

| Algorithm | R package | Hyperparameter values | AUC |
| --- | --- | --- | --- |
| Random Forest | caret | *mtry* = 140<br>*ntree* = 1 | 0.73 |
| Stochastic Gradient Boosting | caret | *interaction.depth* = 33<br>*n.trees* = 1<br>*shrinkage* = 0.9<br>*n.minobsinnode* = 6<br>*bag.fraction* = 0.7 | 0.77 |
| Support Vector Machine with Radial Basis Function Kernel | caret | *sigma* = 0.003362502<br>*C* = 5.51 | 0.57 |
| Random Forest | mikropml | *mtry* = 64<br>*ntree* = 88 | 0.85 |
| Extreme Gradient Boosting | mikropml | *nrounds* = 2<br>*max_depth* = 9<br>*colsample_bytree* = 0.6<br>*eta* = 0.1<br>*gamma* = 0<br>*min_child_weight* = 3<br>*subsample* = 0.7 | 0.73 |
| Support Vector Machine with Radial Basis Function Kernel | mikropml | *sigma* = 0.003362502<br>*C* = 0.0001 | 0.8 |

**Supplementary Table 4.** Genera considered to be important by some ML classifiers trained on CLR-transformed GM data from the study cohort, classified based on their HEI.

| Genus | Group in which it is more abundant | Models that consider it important | Properties and effects on health | References |
| --- | --- | --- | --- | --- |
| *Candidatus Soleaferrea* | Good HEI | SVC (caret)<br>RF (mikropml) | It possesses anti-inflammatory effects by secreting metabolites and intestinal homeostasis protection properties. It is negatively associated with T2D in mice. | [180]–[182] |
| *Roseburia* | Good HEI | SVC (caret)<br>RF (mikropml) | It is inversely associated with HPF consumption and low diet quality. It has been widely reported to decrease its abundance in T2D patients. It possesses anti-inflammatory and antihypertensive properties. | [5], [164], [183] |
| *Bifidobacterium* | Poor HEI | RF (mikropml)<br>SVC (mikropml) | It has been proved to contribute to obesity in mice and humans. Studies have shown a decrease in its abundance in T2D patients. | [184], [185] |
| *Butyricimonas* | Poor HEI | RF (caret)<br>SVC (caret)<br>RF (mikropml)<br>XGBTREE (mikropml) | It is positively associated with normal BMI. Moreover, the *B. virosa* species has been stated to have a beneficial effect on obesity and T2D by promoting SCFA production and thus the secretion of GLP-1. | [158]–[162] |
| *Lachnospiraceae FCS020 group* | Poor HEI | GBM (caret)<br>SVC (caret)<br>RF (mikropml)<br>SVC (mikropml) | It has been associated with protective properties and a healthier diet in mice. | [141] |
| *Sutterella* | Poor HEI | RF (caret)<br>SVC (caret)<br>RF (mikropml) | It is positively associated with higher HPF consumption and a lower diet quality, and is correlated with the likeness of developing T2D. | [120], [163], [186] |

## Notes

https://github.com/victor5lm/tfm

### Competing Interest Statement

The authors have declared no competing interest.

